# Systematic Benchmarking of High-Throughput Subcellular Spatial Transcriptomics Platforms

**DOI:** 10.1101/2024.12.23.630033

**Authors:** Pengfei Ren, Rui Zhang, Yunfeng Wang, Peng Zhang, Ce Luo, Suyan Wang, Xiaohong Li, Zongxu Zhang, Yanping Zhao, Yufeng He, Haorui Zhang, Yufeng Li, Zhidong Gao, Xiuping Zhang, Yahui Zhao, Zhihua Liu, Yuanguang Meng, Zhe Zhang, Zexian Zeng

## Abstract

Recent advancements in spatial transcriptomics technologies have significantly enhanced resolution and throughput, underscoring an urgent need for systematic benchmarking. To address this, we collected clinical samples from three cancer types – colon adenocarcinoma, hepatocellular carcinoma, and ovarian cancer – and generated serial tissue sections for systematic evaluation. Using these uniformly processed samples, we generated spatial transcriptomics data across five high-throughput platforms with subcellular resolution: Stereo-seq v1.3, Visium HD FFPE, Visium HD FF, CosMx 6K, and Xenium 5K. To establish ground truth datasets, we profiled proteins from adjacent tissue sections corresponding to all five platforms using CODEX and performed single-cell RNA sequencing on the same samples. Leveraging manual cell segmentation and detailed annotations, we systematically assessed each platform’s performance across key metrics, including capture sensitivity, specificity, diffusion control, cell segmentation, cell annotation, spatial clustering, and transcript-protein alignment with adjacent CODEX. The uniformly generated, processed, and annotated multi-omics dataset is valuable for advancing computational method development and biological discoveries. The dataset is accessible via SPATCH, a user-friendly web server for visualization and download (http://spatch.pku-genomics.org/).

## Introduction

Spatially resolved transcriptomics integrates high-throughput transcriptomic profiling with spatially contextualized tissue architecture, bridging the gap in single-cell RNA sequencing (scRNA-seq) by linking molecular profiles to their spatial context. With preserved spatial information, this technology offers unprecedented insights into cellular states, tissue organization, and intercellular interactions^1,2,3,4,5^. Its applications span multiple biological disciplines: in neuroscience, it enables high-resolution mapping of neural circuits and molecular connectivity; in developmental biology, it illuminates the molecular mechanisms underlying tissue morphogenesis; and in cancer biology, it provides detailed characterization of tumor microenvironments and immune landscapes^6,7,8^. Driven by its transformative potential, spatial transcriptomics technology has undergone rapid development and innovation.

Spatial transcriptomics (ST) technologies can be broadly categorized into sequencing-based (sST) and imaging-based (iST) platforms, each offering distinct methodologies and advantages. sST platforms enable unbiased whole-transcriptome analysis using poly(dT) oligos to capture target sequences with poly(A) tails on spatially barcoded arrays. These platforms vary in capture efficiency, transcript diffusion control, and spatial resolution, which ranges from microscale to nanoscale. Notable platforms include Visium^9^, DBiT-seq^10^, Patho-DBiT^11^, Stereo-seq^12^, Slide-seq^13^, Slide-seqV2^14^, HDST^15^, sciSpace^16^, PIXEL-seq^17^, Seq-Scope^18^, MAGIC-seq^19^, and Open-ST^20^. Conversely, iST platforms utilize iterative hybridization with fluorescently labeled probes and sequential imaging to achieve single-molecule resolution of gene profiling in situ. iST technologies differ in probe design, signal amplification strategies, imaging modalities, and the target genes to profile. Notable platforms include ISS^21^, CosMx^22^, Xenium^23^, MERFISH^24^, seqFISH^25^, osmFISH^26^, and STARmap^27^. These complementary approaches underscore the strengths of ST, with sST providing comprehensive, unbiased transcriptome-wide coverage, and iST providing high-resolution, targeted gene expression.

Several efforts have been made to benchmark both sST and iST technologies. A comprehensive evaluation by Yue et al.^28^ compared different sST platforms, including Stereo-seq^12^ with 0.5 μm sequencing spot diameter and Visium^9^ with its 55 μm resolution. For iST platforms, Xenium, MERSCOPE, and CosMx were compared using gene panels ranging from 200 to 1000 genes^29,30,31^. A comparative study involving four iST platforms was conducted using in-house and public data, with gene panels of up to 345 genes^32^. Additionally, Austin et al.^33^ evaluated six iST platforms using public datasets, with panel sizes ranging from 99 to 1147 genes. While these studies provide valuable insights, they primarily focus on ST technologies with lower spatial resolution or limited gene panel sizes. Furthermore, many benchmarking studies rely on public datasets generated under varying experimental conditions or with varying tissue types, which often lack consistent ground truth data for robust evaluation. As a result, existing efforts offer only a partial understanding of the latest advancements in ST technologies. This underscores the urgent need for a systematic benchmarking study conducted under unified experimental conditions to comprehensively evaluate the performance and comparative strengths of current high-resolution, high-throughput ST platforms.

Spatial transcriptomics has undergone remarkable advancements, with commercial platforms now achieving subcellular resolution and high-throughput gene detection. Among sST platforms, Stereo-seq v1.3^12^ by BGI employs poly(dT) oligos on the spatially arrayed DNA nanoballs to capture RNA with poly(A) tails at a resolution of 0.5μm. Visium HD^34^ by 10x Genomics utilizes in situ printed poly(dT) oligos to capture probes with poly(A) tails targeting 18,085 genes at a resolution of 2 μm. In parallel, iST platforms, such as CosMx 6K^22^ by NanoString and Xenium^23^ by 10x Genomics, rely on fluorescently labeled probes and sequential imaging to profile 6,175 and 5,001 genes, respectively, offering single-molecule precision. These advancements underscore the pressing need for a systematic benchmark to enable more informed applications and continued innovation in this rapidly evolving field.

In this study, we collected clinical samples from three cancer types and generated serial tissue sections to systematically evaluate five commercially available high-throughput ST platforms with subcellular resolution. To establish ground truth datasets, we profiled proteins in adjacent tissue sections corresponding to each ST platform using CODEX^35^. In parallel, scRNA-seq was performed on the same samples to provide a comparative reference. We manually annotated cell types for both the scRNA-seq and CODEX data, along with cell boundaries in H&E and DAPI-stained images. Leveraging these comprehensive annotations, we systematically evaluated each platform’s performance across critical metrics, including sensitivity, specificity, diffusion control, cell segmentation, cell annotation, spatial clustering, and transcript-protein alignment. The resulting uniformly generated, processed, and annotated multi-omics dataset, comprising 8,133,524 cells, serves as a valuable resource for advancing computational method development and enabling biological discoveries. To facilitate accessibility, we developed a user-friendly web server (SPATCH: http://spatch.pku-genomics.org/) for data visualization, exploration, and download.

## Results

### Sample preparation and profiling for ST benchmarking

To enable a comprehensive and systematic benchmarking of ST platforms, we collected treatment-naïve tumour samples from three patients diagnosed with colon adenocarcinoma (COAD), hepatocellular carcinoma (HCC), and ovarian cancer (OV). To accommodate the sample preparation requirements of each platform, we divided the tumour samples into multiple portions and processed them into formalin-fixed paraffin-embedded (FFPE) blocks, fresh frozen (FF) blocks using optimal cutting temperature (OCT) compound, and disassociated single-cell suspensions (**Fig. 1a**). Serial tissue sections were uniformly generated from each embedded tissue block to ensure consistent evaluation across all platforms. The detailed timelines of sample collection, fixation, embedding, slicing, and gene expression profiling were recorded (**Supplementary Table 1**).

**Fig. 1.**
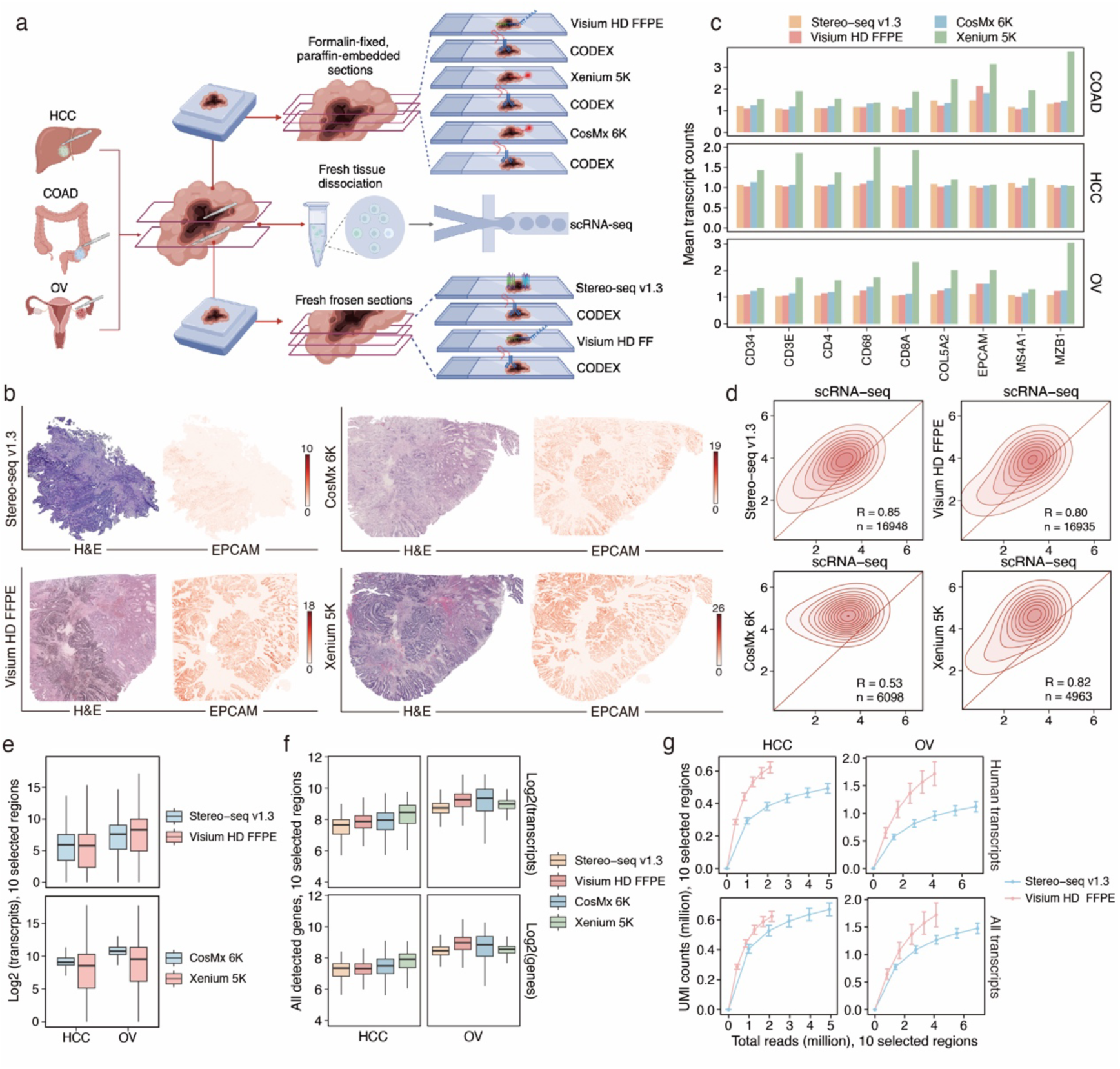
Benchmarking gene detection sensitivity across spatial transcriptomics platforms. **a.** Schematic overview of sample collection, processing, and data generation. Human tumour samples were divided into three sections: one was formalin-fixed and paraffin-embedded (FFPE) for Visium HD FFPE, CosMx 6K, and Xenium 5K profiling; another was embedded in Optimal Cutting Temperature (OCT) for Stereo-seq v1.3 and Visium HD FF profiling; and the remaining tissue was dissociated for single-cell RNA sequencing (scRNA-seq). The spatial distributions of 16 proteins on adjacent sections were profiled with CODEX. **b.** H&E staining and spatial distribution of EPCAM transcripts in COAD sections. Color intensity represents the transcript counts within 8 × 8 μm bins. **c.** Average transcript counts for lineage marker genes across whole sections, calculated at the resolution of 8 μm. **d.** Pearson correlation of gene expression levels between different ST data and scRNA-seq data. For each gene, the total transcript counts across three cancer types were averaged and log10 transformed. The diagonal red line indicates a slope of 1, and color intensity corresponds to relative gene counts. R denotes the correlation coefficient, and n indicates the number of genes included in the analysis. **e.** Log2-transformed total transcript counts of each gene within the ten selected regions (400 × 400 μm) with similar morphology in HCC and OV. **f**. Log2-transformed gene and transcript counts of each 8-μm bin within the ten selected regions with similar morphology in HCC and OV. **g.** Number of human transcripts (top) and all transcripts (bottom) detected by Stereo-seq v1.3 and Visium HD FFPE within the same regions as panel **e**, calculated at stepwise downsampled sequencing depths. Error bars indicate the standard error of the mean (SEM) across the ten selected regions.

We benchmarked five advanced ST platforms – Stereo-seq v1.3, Visium HD FF, Visium HD FFPE, CosMx 6K, and Xenium 5K – selected for their high-throughput gene capture capacity (>5,000 genes), subcellular resolution (≤2um), and wide adoption (commercialized) (**Fig. 1a**). These platforms represent a diverse range of technologies (**Supplementary Table 2**) and utilize overlapping yet distinct gene panels to capture key biological pathways (**Supplementary Fig. 1a, b and Supplementary Table 3, 4**). To establish comprehensive ground truth datasets for robust evaluation, we profiled proteins from adjacent tissue sections corresponding to each ST platform using CODEX and performed scRNA-seq on the same samples (**Fig. 1a**). The uniformly generated reference datasets enabled integrative and cross-modal comparisons across diverse platforms. Of note, the protocol for Visium HD FF became available three months after the initial preparation of samples for other platforms (**Supplementary Table 1**), which may have resulted in RNA degradation in the samples used for Visium HD FF.

### Evaluation of marker gene detection

We first examined the marker gene detection for diverse cell lineages across ST platforms. The epithelial cell marker *EPCAM* showed well-defined spatial patterns across all platforms in COAD, correlating with the hematoxylin and eosin (H&E) staining patterns (**Fig. 1b and Supplementary Fig. 2a**). Xenium 5K demonstrated superior sensitivity for lineage marker gene detection compared to other platforms when analyzed at 8-μm resolution (**Fig. 1c and Supplementary Fig. 2b**). To reduce potential biases from scanning area and tissue morphology, we focused on shared regions among FFPE serial sections (Visium HD FFPE, CosMx 6K, and Xenium 5K). Across these regions, Xenium 5K consistently outperformed other platforms (**Supplementary Fig. 2c**).

To further reduce scanning area difference, we selected ten regions of interest (ROIs, 400 × 400 μm) primarily composed of cancer cells with similar morphology and cell density from each dataset (**Supplementary Fig. 2d**). COAD was excluded from this comparison due to the potential biases caused by the irregular organization of cancer cells. Across the ten ROIs in HCC sections, the capture efficiency for the hepatocyte marker *ALB* varied among Visium HD FFPE, Visium HD FF, and Stereo-seq v1.3 (**Supplementary Fig. 2e**). For the OV sections, the five ST platforms displayed different sensitivities for *EPCAM* detection (**Supplementary Fig. 2f**). Variability in the capture efficiency of Visium HD FF was observed, potentially due to RNA degradation in the samples (**Supplementary Fig. 2e, f**).

### Evaluation of molecular capture efficiency and alignment metrics

We performed correlation analyses for gene expression levels between ST and scRNA-seq data. Stereo-seq v1.3, Visium HD FFPE, and Xenium 5K showed high correlations with scRNA-seq (**Fig. 1d and Supplementary Fig. 3a**). Although CosMx 6K captured more transcripts than Xenium 5K (**Supplementary Fig. 3b**), the total transcript counts revealed a disproportionate elevation across genes showing differing expression levels in scRNA-seq, displaying a skewed pattern (**Fig. 1d**). Similar patterns were observed when restricting the analysis to shared genes (2,522 genes) between the two iST platforms, CosMx 6K and Xenium 5K (**Supplementary Fig. 3c**). This correlation for CosMx 6K remained consistent when using calls detected with varying quality (**Supplementary Fig. 3d**). Cross-platform correlation analyses further highlighted concordance among Stereo-seq v1.3, Visium HD FFPE, and Xenium 5K, underscoring their consistency in describing the varying expression levels of different genes (**Supplementary Fig. 3e**).

We further compared the total transcript counts across genes detected by each ST platform within the ten ROIs. Expression levels should exhibit variation across different gene categories to provide biological insights. Visium HD FFPE and Stereo-seq v1.3 data showed comparable results, while CosMx 6K detected a higher transcript count than Xenium 5K but with less variation between genes (**Fig. 1e**). Similar patterns were observed when using shared genes between CosMx 6K and Xenium 5K (**Supplementary Fig. 4a**). Additionally, consistent patterns were observed when restricting the analysis to shared regions in serial FFPE sections (**Supplementary Fig. 4b**).

In addition to evaluating total transcript counts for individual genes, we analyzed the total number of transcripts and genes detected per 8-μm bin across the ten ROIs, which were roughly comparable across platforms (**Fig. 1f**). Restricting the analysis to shared regions in FFPE serial sections revealed comparable sensitivity, with CosMx 6K showing slightly higher counts than other platforms (**Supplementary Fig. 4c**). When focusing on the common genes shared by the two iST platforms, Xenium 5K detected higher numbers of transcripts and genes per bin compared to CosMx 6K (**Supplementary Fig. 4d**).

For sST platforms, we evaluated quality metrics related to read alignment. Stereo-seq v1.3 had a lower percentage of reads failing UMI quality control compared to Visium HD FFPE and scRNA-seq (**Supplementary Fig. 4e**). However, it showed a higher proportion of reads with barcodes not matching the mask file, indicating a more significant loss of spatial information (**Supplementary Fig. 4e**). Besides, Stereo-seq v1.3 showed a higher proportion of reads mapping to intergenic regions and multiple genomic loci compared to scRNA-seq (**Supplementary Fig. 4f**). To account for sequencing depth variations, we downsampled the reads for Stereo-seq v1.3 and Visium HD FFPE. Visium HD FFPE demonstrated higher sensitivity for detecting human transcriptome at a lower saturation rate (**Fig. 1g**). However, the sensitivity differences between the two platforms were significantly reduced when including all captured transcripts regardless of species (**Fig. 1g**). These findings highlight distinct strengths of each platform in transcript detection and alignment accuracy.

### Evaluation of transcript background noise and diffusion control

Accurate transcript identification is crucial for uncovering the underlying biological mechanisms. In iST platforms, negative probes and codes are used to detect nonspecific binding and fluorescence detection errors. For both iST platforms, signals from negative controls were displayed alongside gene probes targeting human transcriptome (analyzed with 8 × 8 μm bins) (**Fig. 2a**). Overall, CosMx 6K detected more transcripts with a more uniform distribution and exhibited stronger negative control signals compared to Xenium 5K (**Fig. 2b and Supplementary Fig. 5a, b**). Using a spatial autocorrelation metric Moran’s I, we found aggregations of negative control signals in CosMx 6K (**Fig. 2b and Supplementary Fig. 5a, b**). After normalization by the total number of signals, Xenium 5K showed a lower ratio of negative control signals compared to CosMx 6K (**Fig. 2c**). Notably, negative probes consistently exhibited stronger signals than negative codes for both platforms, suggesting nonspecific binding to target transcripts are the primary source of background noise in iST platforms (**Fig. 2b, c and Supplementary Fig. 5a, b**). To mitigate potential biases from low-quality signals, we extracted high-quality calls and re-evaluated the negative control ratio for CosMx 6K, confirming consistent results for negative probes but observing reduced signals for negative codes (**Supplementary Fig. 5c**).

**Fig. 2.**
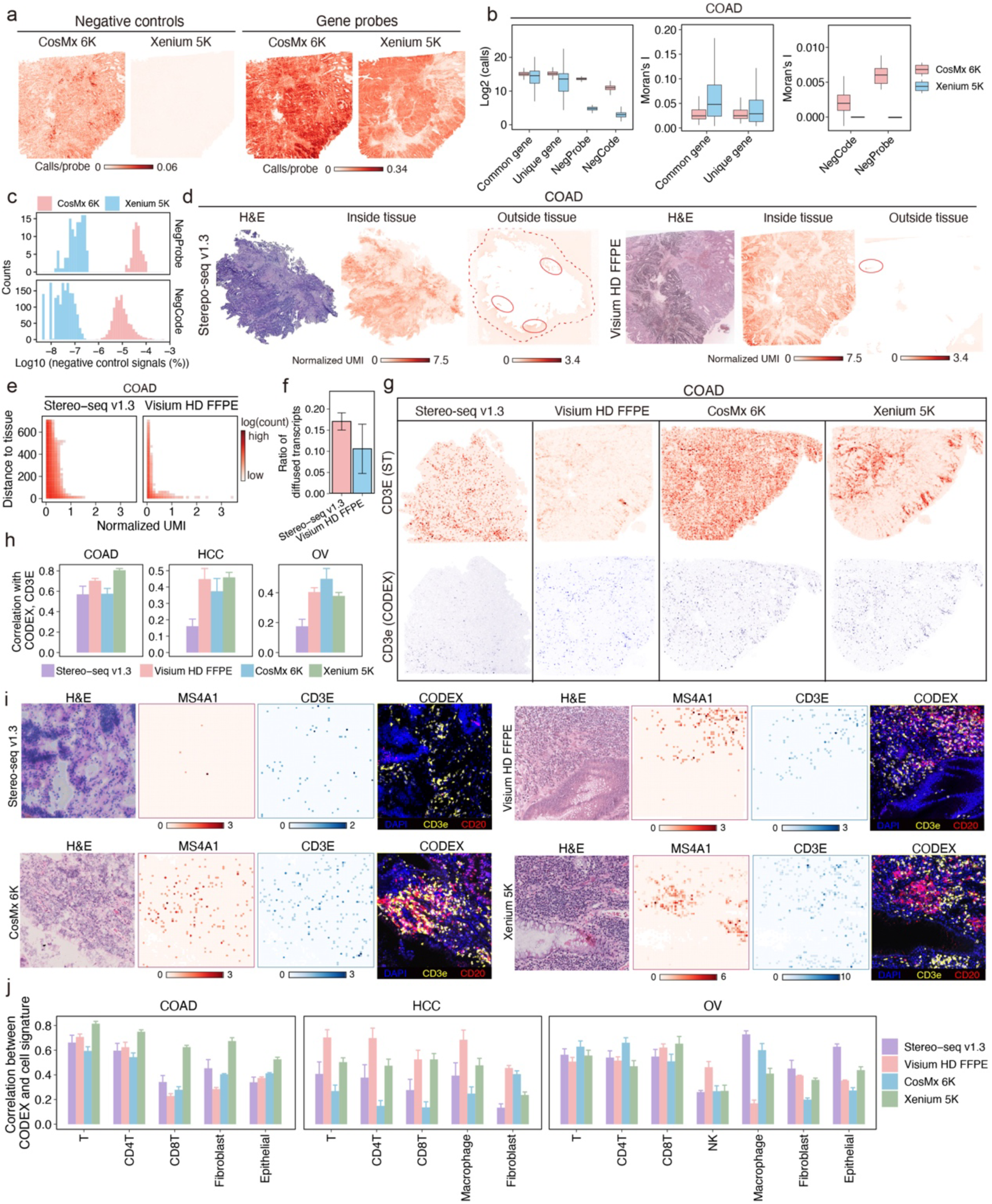
Evaluation of false positives and transcript-protein correlation across spatial transcriptomics platforms. **a.** Spatial distribution of detected negative controls and genes in shared regions between CosMx 6K and Xenium 5K data from COAD sections. Color intensity represents the normalized number of calls in each 8 × 8 μm bin. **b.** Number of calls and Moran’s I for common genes (n = 2552), platform-specific genes (3623 for CosMx 6K and 2449 for Xenium 5K), negative control sequences (NegProbe, 20 for CosMx 6K and 40 for Xenium 5K), and negative control codes (NegCode, 324 for CosMx 6K and 609 for Xenium 5K) detected by CosMx 6K and Xenium 5K in COAD. **c.** Percentage of negative control signals for CosMx 6K and Xenium 5K across three cancer types. **d.** H&E staining and spatial distribution of transcripts detected within and outside the tissue regions of COAD. The red dashed line indicates the Stereo-seq v1.3 region selected for downstream analysis. Solid lines denote regions with high transcript detections outside the tissue. Color intensity indicates mean-normalized transcript count in each 8 × 8 μm bin. **e.** Evaluation of transcript diffusion in COAD. The x-axis represents the mean-normalized transcript count for each bin outside the tissue. The y-axis shows the minimum distance from bins outside the tissue region to the tissue boundary. Color intensity indicates the number of bins. **f.** Relative abundance of transcripts detected within bins outside the tissue region to bins inside the tissue region for Stereo-seq v1.3 and Visium HD FFPE across three cancer types. Error bars represent SEM for the three cancer types. **g.** Spatial distribution of CD3E transcripts (top) from ST data and the corresponding protein (bottom) from adjacent CODEX data in COAD. Data were binned at a 50-μm resolution for visualization. Color intensity denotes transcript counts for ST data and pixel intensity for CODEX data. **h.** Spatial correlation between the abundance of CD3E transcripts and CD3e protein over the spatial grids across three cancer types. Pearson correlation coefficients are reported. Error bars represent SEM for correlations obtained under different grid sizes. **i.** H&E staining and spatial distribution of MS4A1 and CD3E transcripts, along with corresponding protein staining, within regions with high lymphocyte infiltration in COAD. Color intensity denotes transcript counts in 8 × 8 μm bins. **j.** Spatial correlation between CODEX pixel intensity and the corresponding signature scores of ST data for different cell types over the spatial grids. Pearson correlation coefficients are reported. Error bars represent SEM for correlations obtained under different grid sizes.

In sST platforms, the spatial localization of transcripts can be altered during their release onto slides. Both Stereo-seq v1.3 and Visium HD FFPE exhibited transcript diffusion beyond tissue boundaries (**Fig. 2d and Supplementary Fig. 5d**). To evaluate this diffusion, we analyzed transcripts detected outside tissue regions and measured their distances from the tissue boundary (analyzed with 8 × 8 μm bins). To minimize the biases introduced by differences in chip size, we restricted the maximum diffusion distances examined (**Methods**). Quantification of diffusion distances and transcript counts revealed better diffusion control for Visium HD FFPE compared to Stereo-seq v1.3 (**Fig. 2e and Supplementary Fig. 5e**). To account for variations in sequencing depth, we normalized the mean signal of out-tissue bins by the mean signal of in-tissue bins. This analysis revealed a higher proportion of transcripts diffused out of tissue in Stereo-seq v1.3 compared to Visium HD FFPE (**Fig. 2f**).

### Evaluation of spatial concordance with CODEX

We profiled proteins on tissue sections adjacent to each ST section using CODEX, providing a reliable reference for evaluating transcript localization accuracy. The spatial distribution of *CD3E* transcripts detected by ST platforms was compared to the corresponding protein signals in CODEX sections(**Fig. 2g, h**), revealing platform-dependent concordance variations. Analyses of additional markers, such as *CD4*, *CD8A*, and *EPCAM*, further highlighted the discrepancies across ST platforms (**Supplementary Fig. 6a**). Tertiary lymphoid structures (TLS) are aggregates of T cells and B cells that enhance both humoral and cellular immunity, playing a crucial role in anti-tumour responses^36^. In COAD, we identified TLS-like structures and analyzed the spatial distribution of associated transcripts. Concordance between ST and CODEX data for *MS4A1* (B cell marker) and *CD3E* (T cell marker) suggests that ST platforms could recapitulate the cellular organization in TME (**Fig. 2i**).

To address the inherent sparsity of ST data, we focused on cell-level correlations instead of individual markers. Differentially expressed genes for major cell types were identified using paired scRNA-Seq data (**Supplementary Fig. 6b**) and were used to derive cell-type signatures for evaluating spatial concordance with CODEX. Platform– and sample-dependent variations were observed, with Xenium 5K performing better in COAD, Visium HD FFPE showing better concordance in HCC, and CosMx 6K and Stereo-seq v1.3 performing better for certain cell types in OV (**Fig. 2j**). In the selected TLS-like ROIs, we further evaluated cell-type signatures, which demonstrated higher spatial concordance with CODEX than individual markers (**Supplementary Fig. 6c**). Notably, iST platforms exhibited better concordance with CODEX than sST platforms (**Supplementary Fig. 6c**).

In HCC, we leveraged the liver’s complex vascular network, including sinusoids, central veins, and portal triads, to analyze the spatial distribution of *CD34* (endothelial cell marker) near vascular structures. Among the ST platforms, Xenium 5K detected high levels of *CD34* transcripts along vascular edges (**Supplementary Fig. 6d**). In OV, we identified macrophage aggregations and evaluated the distribution of *CD68* transcripts in these regions. All platforms detected substantial *CD68* transcripts, highlighting their effectiveness in detecting macrophage-related markers (**Supplementary Fig. 6e**).

### Evaluation of the single-cell segmentation

Cell segmentation is a critical step for ST platforms with subcellular resolution, heavily impacting downstream analyses. We evaluated the segmentation algorithms implemented within each ST platform with default parameters. Four key metrics describing the cell shapes were included in this comparison: cell size, solidity (where 1 indicates maximal convexity), aspect ratio (length-to-width ratio), and circularity (where 1 represents a perfect circle). Visium HD FFPE was excluded from this analysis due to the unavailability of an official cell segmentation algorithm. Overall, the CosMx 6K segmentation tool produced cells with larger sizes, higher solidity, and better circularity, indicative of more regular and convex cell shapes compared to other platforms (**Supplementary Fig. 7a**-d). To further compare the segmentation accuracy, we selected five regions (500 × 500 μm) from each dataset and manually annotated the boundaries of 72,405 cells to establish ground truth (**Supplementary Fig. 7e**). The comparison between automated segmentation results and manual annotations revealed higher accuracy for iST platforms using DAPI images (**Fig. 3a, b**). To further evaluate segmentation performance, we analyzed the fraction of transcripts located inside the automatically segmented cells. Our results showed that iST platforms had a higher proportion of transcripts confined within the segmented cells compared to the sST platform (**Supplementary Fig. 7f**).

**Fig. 3.**
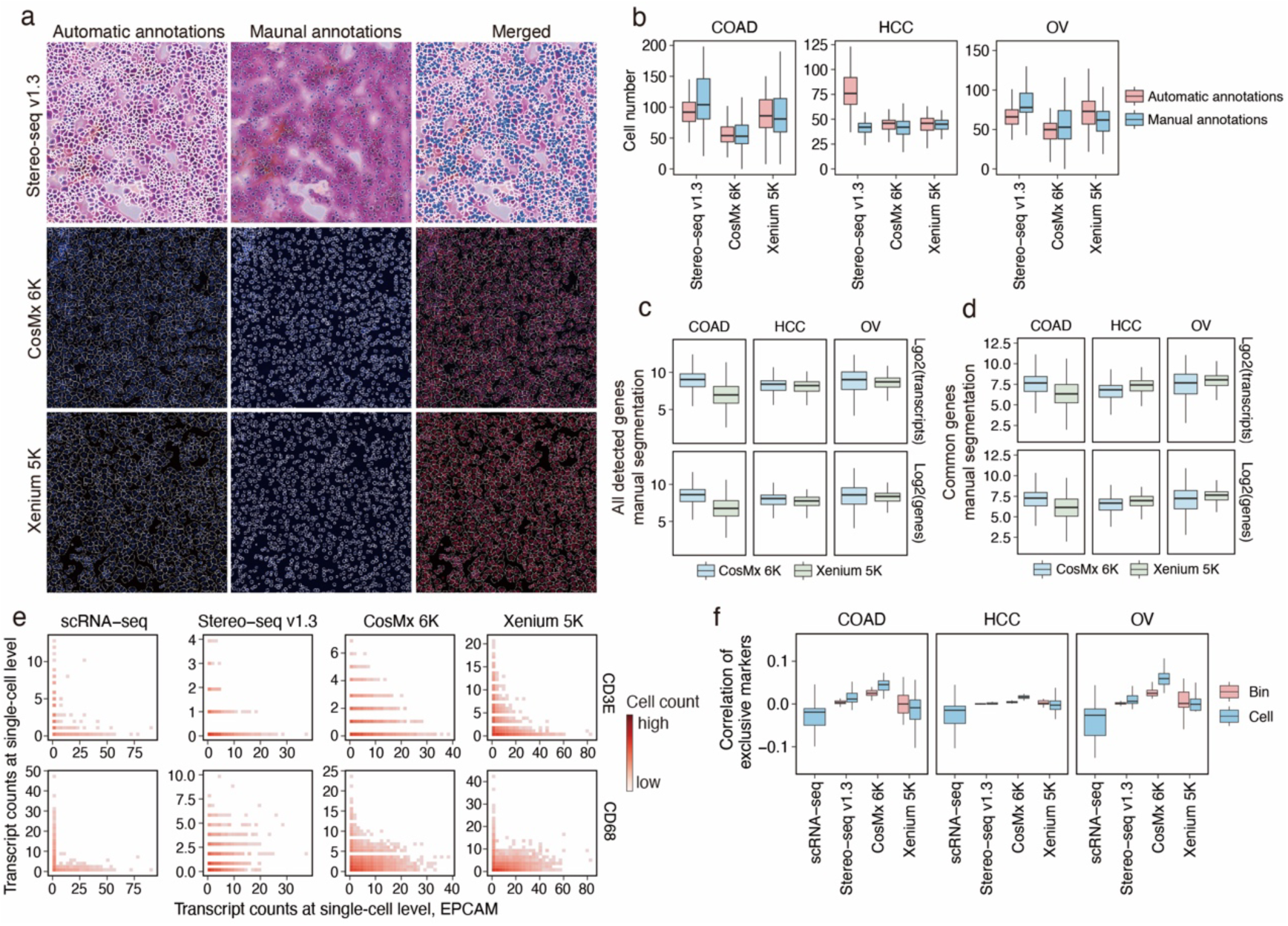
Comparison of cell segmentation. **a.** Cell boundaries generated using automatic cell segmentation algorithms implemented in each ST platform (left) and manual cell segmentation by human annotators (middle). The merged results are shown (right), with white polygons denoting automatic annotations and blue masks indicating manual annotations. **b.** Number of cells annotated by automatic cell segmentation algorithms and human annotators within 125 100 × 100 μm bins. **c-d.** Log2-transformed transcript and gene counts of each manually segmented cell for all detected genes (**c**) or common genes (**d**) detected by the two iST platforms. **e.** Density plot of CD3E/CD68 and EPCAM expressions in COAD. Only cells containing at least one transcript of either CD3E/CD68 or EPCAM were included. Color intensity indicates the number of single cells. **f.** Expression correlation of marker genes expected to be exclusively expressed in different major lineages (n = 36 gene pairs).

Following single-cell segmentation, transcript and gene counts per cell were compared across platforms. Our analysis showed that scRNA-seq consistently detected more transcripts and genes per cell compared to the ST platforms (**Supplementary Fig. 7g)**. Marker gene expression levels were also higher in scRNA-seq data compared to ST data (**Supplementary Fig. 7h**). However, when focusing on the common gene set (2,522 genes) shared among the two iST platforms and scRNA-seq, the iST platforms demonstrated comparable performance to scRNA-seq (**Supplementary Fig. 7i**). Among the ST platforms, CosMx 6K detected the highest number of transcripts and genes per cell (**Supplementary Fig. 7g**). To reduce potential biases introduced by segmentation tools, we also evaluated transcript and gene counts using manually segmented cells within the paired regions between CosMx 6K and Xenium 5K data. This analysis confirmed that CosMx 6K captured more transcripts and genes per cell compared to Xenium 5K (**Fig. 3c**). When restricted to common genes, CosMx 6K outperformed Xenium 5K in COAD (**Fig. 3d**).

A critical criterion for accurate single-cell segmentation is the assignment of mutually exclusive genes into different cells. At the single-cell level, substantial co-expression between markers of epithelial cells (*EPCAM*) and immune cells (*CD3E*/*CD68*) was observed across ST data in COAD, with levels notably higher than those detected by scRNA-seq (**Fig. 3e**). Additionally, by comparing the co-expression at the single-cell level (**Fig. 3e**) to bin level (**Supplementary Fig. 7j**), segmentation algorithms from different ST platforms showed varying ability in correctly assigning *EPCAM*, *CD68*, and *CD3E* into different cells. To further evaluate the co-expression patterns, we calculated expression correlations for 36 gene pairs derived from combinations of nine marker genes (**Supplementary Table 5**) that are exclusively expressed in different cell lineages. ScRNA-seq data consistently exhibited lower co-expression levels than ST data (**Fig. 3f**). Among the ST platforms, Xenium 5K showed better separation of marker gene pairs after segmentation (**Fig. 3f**).

### Evaluation of cell clustering and cell type annotation

Cell clustering is pivotal in describing various cell states and is widely used in discovering novel cell types under both normal and disease conditions. We performed cell clustering based on gene expression at the single-cell level (8-μm bin level for Visium HD FFPE) to assess clustering performance (**Fig. 4a**). Clustering quality, as measured by the average silhouette score, revealed that scRNA-seq achieved superior clustering performance compared to ST platforms (**Fig. 4b**). Among the ST platforms, iST technologies demonstrated better clustering quality than sST platforms, highlighting the advantage of their higher spatial resolution (**Fig. 4b**).

**Fig. 4.**
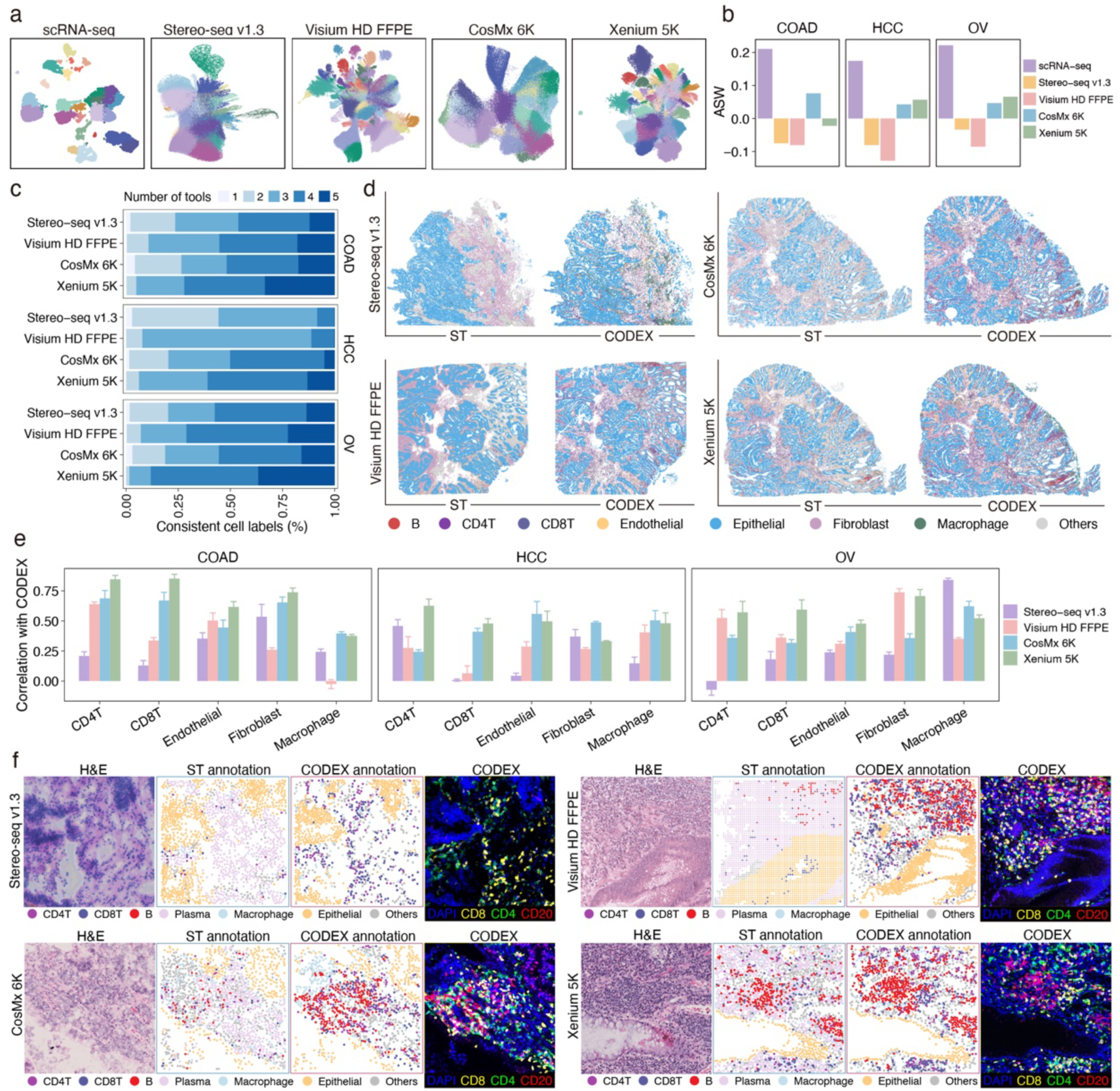
Comparative analysis of cell type annotations, immune cell detection, and spatial alignment with adjacent CODEX. **a.** UMAP representation of ST and scRNA-seq data from COAD. Each point represents a single cell for scRNA-seq, Stereo-seq v1.3, CosMx 6K, and Xenium 5K data. For Visium HD FFPE, each point corresponds to an 8 × 8 μm bin. Distinct colors denote different clusters. **b.** Average silhouette width (ASW) of clustering results across platforms, with higher scores indicating better clustering quality. **c.** Proportion of cells consistently annotated as the same cell type by multiple automatic annotation tools. Colors represent the number of tools that have consistent annotations. **d.** Comparison of cell type annotations derived from ST data with those from adjacent CODEX. **e.** Correlation of cell counts between ST data and adjacent CODEX for each cell type over spatial grids. Pearson correlation coefficients are shown. Error bars represent SEM for correlations obtained under different grid sizes. **f.** Spatial distribution of major cell types within regions of high lymphocyte infiltration (500 × 500 μm). The H&E staining, cell type annotations of ST and CODEX data, and protein staining for CD8, CD4, and CD20 are shown together.

Cell type annotation is another critical step in downstream analysis. To evaluate the ability of different ST platforms to detect diverse cell types, we transferred annotations from scRNA-seq data to ST data using five annotation tools, including SELINA^37^, Celltypist^38^, Spoint^39^, Tangram^40^, and TACCO^41^. Among the evaluated ST platforms, CosMx 6K and Xenium 5K recovered more cell types (**Supplementary Fig. 8a**). Additionally, Xenium 5K showed a higher proportion of cells consistently annotated as the same cell type across tools, indicating robust annotation reliability (**Fig. 4c**). Annotation discrepancy across tools was further assessed by calculating entropy, with higher entropy indicating more consistent cell type composition revealed by different tools. Xenium 5K performed well in COAD and OV, while CosMx 6K showed better performance in HCC (**Supplementary Fig. 8b**). To derive final cell type annotations, results from the five tools were integrated using a majority vote approach (**Methods**). After cell type annotation, we evaluated the alignment of ST data with scRNA-seq data by performing pairwise correlation analyses of gene expression for each cell type. Stereo-seq v1.3 exhibited good performance in COAD and OV, while Xenium 5K performed well in HCC and OV (**Supplementary Fig. 8c**). We further visualized marker gene expression across the annotated cell types. Xenium 5K exhibited the most distinct expression patterns (**Supplementary Fig. 8d**), facilitating more accurate cell type annotations.

We examined the cell type annotation accuracy of each ST data by comparing it with the adjacent CODEX data. In COAD, all ST data exhibited high concordance with CODEX in terms of cellular organization, with iST data showing better alignment (**Fig. 4d**). To evaluate the ability of ST platforms to detect immune cells, we quantified the total numbers of CD4+ T cells, CD8+ T cells, and macrophages in both ST and CODEX data (**Supplementary Fig. 9a**). Platforms with higher concordance with CODEX were considered more effective in immune cell detection. Overall, iST platforms outperformed sST platforms in identifying lymphocytes characterized by smaller cell sizes (**Supplementary Fig. 9a**). Additionally, we performed correlation analyses of cell counts across spatial grids between ST data and adjacent CODEX data for each cell type. These analyses revealed platform-specific strengths and limitations in cell type identification, with varying levels of consistency between ST platforms and CODEX (**Fig. 4e and Supplementary Fig. 9b**).

### Evaluation of immune cell detection

We assessed the ability of different ST platforms to discriminate diverse immune cell populations in lymphocyte-enriched regions. Lymphocyte aggregates were identified in COAD sections for each ST platform (**Fig. 4f**), with marker genes associated with CD4+ T cells (*CD4*), CD8+ T cells (*CD8A*), B cells (*MS4A1*), plasma cells (*MZB1* and *CD38*) effectively captured (**Supplementary Fig. 9c**-f). Xenium 5K demonstrated high efficiency and accuracy in identifying CD4+ T cells, CD8+ T cells, and B cells (**Fig. 4f**).

Although Visium HD FFPE exhibited high sensitivity and specificity for markers such as *CD3E*, *CD4*, *CD8A*, and *MS4A1* (**Fig. 2i and Supplementary Fig. 9c, d**), it demonstrated limited ability to distinguish CD4+ T cells, CD8+ T cells, and B cells from plasma cells (**Fig. 4f and Supplementary Fig. 9e, f**). This discrepancy is likely due to bin-level analyses resulting in mixed transcripts from neighboring cell types. Beyond COAD, we evaluated endothelial cell distribution in HCC. Both CosMx 6K and Xenium 5K successfully identified endothelial cells distributed along blood vessels, in agreement with anatomical expectations (**Supplementary Fig. 9g**). In OV, macrophage-enriched regions were detected by all ST platforms, demonstrating their robust capacity to identify macrophages (**Supplementary Fig. 9h**).

Advancements in iST platforms, particularly their ability to capture large gene panels, offer the potential for improved characterization of cell subtypes. To evaluate each platform’s ability to recover T cell subtypes, we first established a reference using matched scRNA-seq data (**Supplementary Fig. 10a**). Annotations were transferred to the ST data using five annotation tools, and performance was benchmarked by assessing the consistency of cell type assignments. Among the platforms, CosMx 6K and Xenium 5K successfully recovered the highest number of cell subtypes (**Supplementary Fig. 10b**). Additionally, Xenium 5K demonstrated a higher proportion of cells consistently annotated as the same cell type across tools, indicating high annotation reliability (**Supplementary Fig. 10c**).

### Evaluation of spatial clustering and spatial patterns

Spatial clustering plays a key role in identifying functional cellular aggregates within tissues. To evaluate the extent to which ST data can recapitulate spatial patterns observed in CODEX, we performed spatial clustering on both datasets using CellCharter^42^ (**Methods**). Concordance between clustering results from ST and CODEX data served as a performance measure, with higher concordance indicating better clustering fidelity (**Fig. 5a and Supplementary Fig. 11a**-b). In COAD, Visium HD FFPE and Xenium 5K showed higher concordance with CODEX. CosMx 6K exhibited the highest similarity to CODEX in HCC. The performance of different ST platforms was roughly comparable in OV (**Fig. 5b and Supplementary Fig. 11c**).

**Fig. 5.**
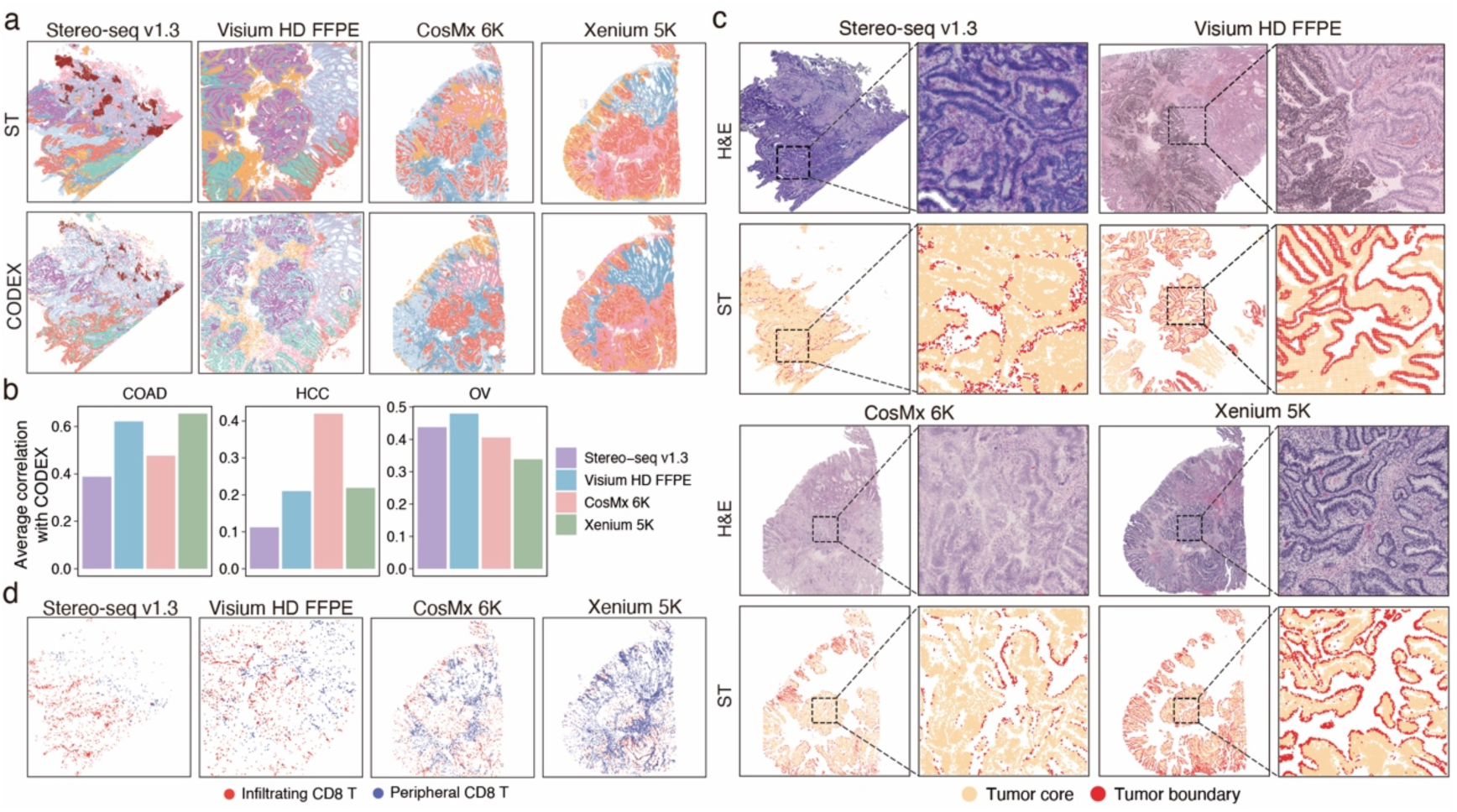
Comparison of spatial clustering and cell distributions. **a.** Spatial clustering of ST and CODEX data from COAD sections, with distinct colors denoting different spatial clusters. **b.** Correlation of cluster proportions between ST and CODEX data across 100 × 100 μm spatial grids. Pearson correlation coefficients are reported. **c.** H&E staining and localization of malignant cells within the tumour core and at the tumour boundary. **d.** Spatial distribution of tumour-infiltrating CD8+ T cells and peripheral CD8+ T cells within the COAD sections.

Spatial clustering also enabled the identification of spatially distinct subtypes, which may exhibit transcriptomic variations linked to tissue architecture. All ST platforms successfully identified clusters preferentially localized at the tumour boundary or within the tumour core, although the delineation of these boundaries varied among platforms (**Fig. 5c, Supplementary Fig. 11d**). Additionally, CD8+ T cells were categorized into infiltrating CD8+ T and peripheral CD8+ T cells based on their co-localization with cancer cells, demonstrating the utility of spatial clustering for characterizing immune cell localization (**Fig. 5d**).

## Discussion

In this study, we comprehensively benchmarked five commercialized high-throughput ST platforms with subcellular resolution. To ensure reliable assessments, we uniformly processed clinical samples to generate high-quality and comparable datasets. Matched scRNA-seq and proteomic data, along with manually annotated cell boundaries, provided robust reference baselines. Using these datasets and annotations, we systematically assessed each ST platform across key performance metrics, including sensitivity, specificity, diffusion control, cell segmentation accuracy, cell annotation reliability, spatial clustering, and transcript-protein alignment. Given the inherent complexity and heterogeneity of the TME, this comprehensive evaluation provides critical insights and practical guidance for selecting the most suitable ST platform tailored to specific research and clinical applications.

sST platforms provide transcriptome-wide profiling capabilities, each with unique advantages. Stereo-seq v1.3, with poly(dT)-based capture method, enables unbiased transcriptome exploration, detecting a wide range of transcripts, including noncoding RNAs, bacterial RNAs, and viral RNAs. This versatility is ideal for studying diverse biological systems, though requiring higher sequencing depth to achieve comparable coverage to targeted platforms such as Visium HD FFPE. Visium HD FFPE, on the other hand, uses a panel of predesigned probes to target 18,085 human genes or 19,405 mouse genes. By capturing shorter probes rather than complete mRNA molecules, Visium HD FFPE achieves enhanced diffusion control, minimizes spatial artifacts, and demonstrates higher concordance with protein-based datasets like CODEX, making it well-suited for applications requiring high spatial precision.

High-throughput iST platforms, such as CosMx 6K and Xenium 5K, provide unique advantages and face challenges compared to sST technologies. By employing fluorescent probes and high-resolution imaging techniques, iST platforms achieve single-molecule resolution, enabling precise transcript localization within individual cells. Furthermore, iST platforms detect transcripts in situ, effectively avoiding diffusion and enhancing spatial mapping accuracy. This high spatial resolution and accuracy is particularly beneficial for analyzing complex tissue architectures and identifying small cell types, such as lymphocytes, where spatial fidelity is crucial. However, their reliance on predefined gene panels limits their ability to perform unbiased, transcriptome-wide profiling. Although CosMx 6K and Xenium 5K have expanded gene panels (6,175 and 5,001 genes, respectively), they are less comprehensive than sST technologies, which can capture the entire transcriptome. Thus, while iST platforms excel in spatial precision, their restricted gene coverage may limit exploratory transcriptomic analyses.

Accurate cell segmentation is essential for high-resolution spatial transcriptomics, as it directly affects downstream analyses, including cell type annotation and spatial clustering. In our study, platforms utilizing DAPI staining demonstrated superior performance compared to those based on H&E staining. This difference is likely due to artifacts in H&E images, such as impurities and air bubbles, which can be misidentified as cells, as well as the less distinct nuclear boundaries inherent to H&E staining. High-throughput iST platforms, such as CosMx 6K and Xenium 5K, improve segmentation accuracy by integrating multiplexed immunofluorescence data, leveraging both nuclear and protein localization signals. Despite exhibiting more irregular segmentation shapes, Xenium 5K effectively distinguished different cell types and minimized transcript mixing between adjacent cells. Conversely, Visium HD FFPE, with its 2 µm resolution, provides subcellular detail but presents challenges for single-cell segmentation, as individual cells may be divided across multiple spots. These findings highlight the need for robust imaging techniques and optimized computational algorithms to achieve reliable cell segmentation in ST platforms.

Our study has several limitations that should be taken into account. First, we focused on commercialized ST platforms with larger gene panels, excluding those with smaller panels or limited commercial availability, which may narrow the scope of our comparisons. Second, CODEX was performed on adjacent tissue sections instead of the original ST sections, which provided essential reference but also introduced morphological discrepancies that may impact direct comparisons across platforms. Additionally, because Stereo-seq v1.3 relies on fresh frozen sections, its comparison with FFPE tissues was inherently constrained by structural differences associated with sample preparation methods. Third, our alignment and segmentation analyses utilized commercial pipelines, representing standardized workflows commonly used by researchers but potentially missing insights from custom computational optimizations. Lastly, as our study exclusively used freshly collected samples, the generalizability of our findings to archived or long-term stored samples remains unclear, underscoring the need for further validation in a broader range of sample types and conditions.

Despite its limitations, this study makes a critical contribution to the field of spatial transcriptomics by comprehensively benchmarking commercially available high-throughput, subcellular-resolution ST platforms. The resulting multi-omics dataset— uniformly generated, processed, and annotated—encompasses spatial transcriptomics, matched CODEX and scRNA-seq data, and detailed manual annotations of cell types and boundaries. Accessible via SPATCH (https://spatch.pku-genomics.org<u>/</u>), this resource addresses a critical gap in high-quality ground truth data for spatial transcriptomics and offers a robust foundation for advancing computational methods and driving continued innovation in the field.

Looking ahead, iST platforms are poised to expand their coverage to encompass unbiased transcriptome-wide profiling across diverse tissues. Concurrently, sST platforms are expected to achieve higher resolution, improved diffusion control, and increased capture sensitivity. Both platform types are anticipated to advance toward seamless integration with proteomics, epigenomics, and other modalities to achieve multi-omics profiling. These developments will provide an unprecedented view of cellular interactions and molecular mechanisms within complex tissue environments, fueling further discoveries and biological insights.

## Methods

### Human sample collection and preprocessing

This study was approved by the Research and Biomedical Ethical Committee of Peking University (IRB00001052-24061) and conducted following pertinent ethical regulations. All patients provided informed consent for collecting clinical information and tumour samples. All protocols adhered to the Interim Measures for the Administration of Human Genetic Resources, administered by the Ministry of Science and Technology of China. The tumour specimens were obtained from 3 patients, with each individual presenting a distinct cancer diagnosis: COAD, HCC, and OV, at the Chinese PLA General Hospital and Peking University People’s Hospital. Necrotic areas and regions adjacent to major blood vessels were excluded during collection. Each tissue was further evenly divided into three sections. The middle portion was submerged in the MACS^®^ Tissue Storage Solution (Miltenyi #130-100-008) and further processed for scRNA-seq. One of the remaining sections was fixed in a 10% neutral formalin fixing solution (Solarbio #G2161) for 24 to 48 hours before paraffin embedding. The other section was embedded in 4°C OCT compound (Sakura #4583), quickly frozen on dry ice, and transferred to a –80 °C freezer for storage until further experimentation. The entire process was completed within 30 minutes to minimize RNA degradation. Serial sections of FFPE samples were prepared at Peking University and loaded onto platform-specific chips under the supervision of trained technicians for Visium HD FFPE, Xenium 5K, and CosMx 6K. Sections adjacent to all ST sections were reserved for subsequent CODEX profiling.

### Stereo-seq v1.3 data generation

Stereo-seq v1.3 assay is compatible with OCT-embedded tissues and H&E staining. Sample preparation and sectioning followed the Guide for Fresh Frozen Samples on Stereo-seq Chip Slides (Document No.: STUM-SP001). RNA quality was evaluated to determine whether to proceed with the following experiments. Only samples with RIN value ≥ 6 are acceptable for further procedures. Cryosections were cut at a thickness of 10 μm in a Leika CM1950 cryostat. H&E staining, in situ reverse transcription, amplification, library construction, and sequencing followed the User Manual of the Stereo-seq Transcriptomics Set v1.3 (STOmics, #201ST13114 or 211ST13114). Tissue sections were loaded onto the Stereo-seq chip (generated by BGI, China; with a maximum size of 1 x 1 cm) and fixed in pre-cooled methanol. Hematoxylin-eosin staining was performed prior to tissue permeabilization. RNA was released from the permeabilized tissue, captured by the DNA nanoball (DNB), and subsequently underwent in situ reverse transcription. Following reverse transcription, tissue sections were removed to release complementary DNA (cDNA), which was purified using the VAHTSTM DNA Clean Beads (VAZYME #N411-02) and Stereo-seq 16 Barcode Library Preparation Kit (STOmics, #101KL160 or 111KL160). For library construction, 100 ng of cDNA was utilized for fragmentation and amplification. PCR products were purified using the VAHTSTM DNA Clean Beads. Ultimately, the purified PCR products were used for DNB production, and the libraries were sequenced using the MGI DNBSEQ-T7 sequencer.

### Visium HD for FFPE data generation

Visium HD assay is compatible with FFPE-embedded tissues and H&E staining. RNA quality of FFPE samples was assessed by calculating the percentage of RNA fragments >200 nucleotides (DV200) extracted from tissue sections. DAPI and H&E staining were also used to assess tissue morphology before performing the Visium HD assay. Tissue sections were cut at a thickness of 5 μm following the Visium HD FFPE Tissue Preparation Handbook (CG000684, 10x Genomics), spread out in RNA enzyme-free water at 42℃, and loaded onto the slides prepared in advance (Fisher Scientific #1255015). Subsequently, these slides were air-dried at room temperature for 30 minutes and baked at 42℃ for 3 hours. The subsequent experiments were carried out after drying overnight at room temperature.

Tissue sections were subjected to deparaffinization, H&E staining, and imaging following the Visium HD FFPE Tissue Preparation Handbook (CG000684, 10x Genomics). Probe hybridization, probe ligation, Visium HD slide preparation, probe release, extension, library construction, and sequencing followed the Visium HD Spatial Gene Expression Reagent Kits User Guide (CG000685, 10x Genomics). The tissue sections were destained and decrosslinked after H&E staining. The human whole transcriptome probe panel, consisting of about three specific probes for each target gene, was added to the tissue sections. After hybridization, the Probe Ligation Enzyme (PN-2000425, 10x Genomics) was added to establish connections between the probe pairs hybridized to RNA, resulting in the formation of ligation products. The subsequent release and capture of these probes within the 6.5 x 6.5 mm capture areas were facilitated by the Visium CytAssist instrument following the User Guide. Treatment with RNase Enzyme and Perm Enzyme detached the single-stranded ligation products from the tissue and directed them onto the Visium HD Slide for capture. These ligation products were then elongated by adding the Spatial Barcode, UMI, and partial Read1 primer. Subsequent elution and amplification of the ligation products prepared them for indexing through the sample index PCR. The final libraries were cleaned up by SPRIselect. Sequencing was performed on an Illumina NovaSeq 6000 to obtain paired-end reads.

### Visium HD for FF data generation

Visium HD assay is also compatible with OCT-embedded tissues and H&E staining. RNA quality of the FF samples was assessed by calculating the RIN of RNA extracted from tissue sections. Tissue sections with RIN ≥ 6 are optimal for the Visium HD assay. DAPI and H&E staining were also used to assess tissue morphology before performing the Visium HD assay. Tissue sections were cut at a thickness of 10 μm in a Leika CM1950 cryostat following the Visium HD Fresh Frozen Tissue Preparation Handbook (CG000763, 10x Genomics) and loaded onto the slides prepared in advance (Fisher Scientific #1255015). Tissue sections were subjected to fixation, H&E staining, and imaging following the Visium HD FF Tissue Preparation Handbook (CG000763, 10x Genomics). After destaining and permeabilization, probe hybridization, probe ligation, Visium HD slide preparation, probe release, extension, library construction, and sequencing were performed following the Visium HD Spatial Gene Expression Reagent Kits User Guide (CG000685, 10x Genomics). The release and capture of the probes within the 6.5 x 6.5 mm capture areas were facilitated by the Visium CytAssist instrument following the User Guide. The final libraries were sequenced on an Illumina NovaSeq 6000 to obtain paired-end reads.

### Xenium 5K data generation

Xenium 5K assay is compatible with FFPE-embedded tissues and H&E staining. RNA quality of the tissue block was assessed by calculating the percentage of RNA fragments >200 nucleotides (DV200) extracted from tissue sections. DAPI and H&E staining were also used to assess tissue morphology before performing the Xenium 5K assay. Tissue sections were cut at a thickness of 5 μm following the Xenium In Situ for FFPE-Tissue Preparation Guide (CG000578), spread out in RNA enzyme-free water at 42℃, and attached to the Xenium slides (PN-3000941, 10x Genomics) within the sample area (with a maximum size of 10.45 x 22.45 mm) without overlapping with the surrounding fiducials. The slides were dried at room temperature for 30 minutes and baked at 42℃ for 3 hours. The follow-up experiment was carried out after drying overnight at room temperature.

After drying overnight, the Xenium slides were subjected to deparaffinization and decrosslinking following the Xenium In Situ Protocol for FFPE-Deparaffinization and Decrosslinking (CG000580, 10x Genomics). Priming hybridization, RNase treatment & polishing, probe hybridization, probe ligation, amplification, cell segmentation staining, autofluorescence quenching, and nuclear staining followed the Xenium Prime In Situ Gene Expression with optional Cell Segmentation Staining (CG000760, 10x Genomics). The Xenium Prime 5K Human Pan Tissue & Pathways Panel (PN-1000724, 10x Genomics) targeting 5001 individual human genes was used for the assay. After priming hybridization, RNase treatment & polishing steps, the Xenium slides were incubated with probes at 50℃ for 16-24 hours for probe hybridization and then washed with PBS-T. Then the slides were subjected to probe ligation at 42℃ for 30 minutes, amplification enhancement at 4℃ for 2 hours, and amplification at 30℃ for 1.5 hours. Following additional washing procedures, the slides underwent cell segmentation staining, then treatment with an autofluorescence suppressor and nuclear staining. The slides were loaded onto Xenium Analyzer (PN-1000529, 10x Genomics) according to the Xenium Analyzer User Guide (CG000584, 10x Genomics) and ran for about 90 hours. The Xenium Onboard Analysis pipeline v.3.1.0 (10x Genomics) was run directly on the instrument for imaging processing, cell segmentation, image registration, decoding, deduplication, and secondary analysis. After that, the slides were washed to perform post-run H&E staining.

### CosMx 6K data generation

CosMx 6K assay is compatible with FFPE-embedded tissues and H&E staining. RNA quality of the tissue block was assessed by calculating the percentage of RNA fragments >200 nucleotides (DV200) extracted from tissue sections. DAPI and H&E staining were also used to assess tissue morphology before performing the CosMx 6K assay. Tissue sections were cut at a thickness of 5 μm following the CosMx SMI Manual Slide Preparation for RNA Assays (MAN-10184-02, NanoString Technologies), spread out in RNA enzyme-free water at 42℃, and attached to the slides (CIITOTEST #188105) within the scan area (with a maximum size of 2.0 x 1.5 cm). The slides were dried at room temperature for 30 minutes and baked at 65℃ for 30 minutes. The follow-up experiment was carried out after drying overnight at room temperature.

Deparaffinization, target retrieval, protease digestion, blocking, hybridization, stringent washing, blocking, nuclear and segmentation markers staining, and imaging followed the CosMx SMI Manual Slide Preparation for RNA Assays (MAN-10184-02, NanoString Technologies). Human 6K Discovery Panel, 6K-plex, RNA (#121500041, NanoString Technologies) was used. The tissue sections were deparaffinized, subjected to target retrieval at 100℃ for 15 minutes, treated with protease for digestion at 40℃ for 30 minutes, incubated with applied fiducials for 5 minutes, post-fixed, blocked, and incubated with the human 6K Discovery Panel overnight. The slides were washed and blocked, followed by nuclear staining. Then the sections were incubated with Marker Stain Mix (PanCK, CD45) and Cell Segmentation Mix (CD298, B2M) using CosMx^TM^ Human Universal Cell Segmentation Kit (RNA) (121500020, NanoString Technologies). The slides were washed again and loaded onto the CosMx SMI system (cat #101000, S/N: SMI_2307H0124) for UV bleaching, imaging acquisition, cycling processing, and scanning according to the Instrument User Manual (MAN-10161-05, NanoString Technologies). The raw images were subsequently decoded using Atomx (v.1.3.2). Finally, the slides were washed to perform post-run H&E staining.

### CODEX data generation

Tissues embedded in FFPE were sliced, spread out in RNA enzyme-free water at 42℃, and loaded onto the slides (Fisher Scientific #1255015) within the scan area (with a maximum size of 3.5 x 1.8 cm). The slides were dried at room temperature for 30 minutes and baked at 65℃ for 30 minutes. The follow-up experiment was carried out after baking overnight at 60℃. A 16-plex commercial antibody panel was used to target 16 proteins. The sample preparation, tissue staining, and imaging followed the PhenoCycler-Fusion User Guide_2.2.0 (PD-000011 REV M, Akoya Biosciences). After overnight baking, the slides were subjected to deparaffinization and antigen retrieval using a sodium citrate solution for 20 minutes at 11.6PSI/110°C. Subsequently, the slides were washed using a hydration buffer (P/N 7000017, Akoya Biosciences) and incubated in a staining buffer (P/N 7000017, Akoya Biosciences) at room temperature for 20 minutes. A mixture of antibodies, blocking solution, and staining solution was prepared. The slides were incubated with the staining mix at room temperature for 3 hours. The antibody dilution ratio was determined based on pre-tests, along with the cycle information summarized in **Supplementary Table 6**. Following staining, the tissues were sequentially fixed with PFA, ice-cold methanol, and a final fixative solution. The slides were washed, loaded onto the PhenoCycler-Fusion instrument (PhenoCycler-Fusion 2.0), and imaged according to the instrument’s instructions.

For the FF samples, cryosections were cut at a thickness of 10 μm in a Leika CM1950 cryostat, loaded onto the slides (Fisher Scientific #1255015) within the scan area (with a maximum size of 3.5 x 1.8 cm) and stored at –80℃ before the experiment. The sample preparation, tissue staining, and imaging followed the PhenoCycler-Fusion User Guide_2.2.0 (PD-000011 REV M, Akoya Biosciences). The slides were dried and warmed for 5 minutes at room temperature, fixed in acetone for 10 minutes, incubated with the hydration buffer (P/N 7000017, Akoya Biosciences), fixed with 1.6% PFA for 10 minutes, and finally balanced with staining buffer (P/N 7000017, Akoya Biosciences) at room temperature for 20 minutes. A mixture of antibody, blocking solution, and staining solution was prepared. Then the slides were incubated with the staining mix at room temperature for 3 hours. The antibody dilution ratio was consistent with the staining of FFPE samples. Following staining, the tissues were sequentially fixed with PFA, ice-cold methanol, and a final fixative solution. The slides were washed, loaded onto the PhenoCycler-Fusion instrument (PhenoCycler-Fusion 2.0), and imaged according to the instrument’s instructions.

### scRNA-seq data generation

Single-cell suspensions from primary human tumour tissue were generated using the Tumour Dissociation Kit (Miltenyi Biotec, #130-095-929). The proportions of living cells were more than 85% for all samples. The single-cell suspension was processed with the Chromium Single Cell 3’ GEM, Library & Gel Bead Kit v3.1 (10x Genomics, PN-1000268) and loaded onto a Chromium Single Cell Chip (Chromium Single Cell G Chip Kit, 10x Genomics, PN-1000120) according to the manufacturer’s instructions for co-encapsulation with barcoded Gel Beads. The captured cells were lysed, and the released RNA was barcoded through reverse transcription in individual single-cell gel beads in the emulsion (GEMS). In each droplet, cDNA was generated and amplified through reverse transcription on a T100 PCR Thermal Cycler (Bio-Rad) at 53℃ for 45 minutes, followed by 85℃ for 5 minutes and a hold at 4℃. Then, cDNA concentration and quality were assessed using a Qubit Fluorometer (Thermo Scientific) and bioanalyzer 2100 (Agilent), respectively. scRNA-seq libraries were then constructed and sequenced on the Illumina platform according to the manufacturer’s introduction.

### Collection of gene sets for cytokines, chemokines, ligands, receptors, membrane proteins, and transcription factors

Cell membrane proteins, which span or anchor/embed within the plasma membrane, facilitate communication between cells and the extracellular environment. Both experimental and computational approaches have been employed to identify and predict cell-surface membrane proteins. However, each method has inherent limitations, often resulting in incomplete coverage and false positives^43,44,45^. Among the various resources available, we selected the latest and most comprehensive database related to cancer research^46^. Ligands, receptors, cytokines, and transcription factors were also collected from previously published studies and databases^47,48,49,50,51^.

### Data preprocessing

scRNA-seq data were processed with cellranger (v.7.0.0). Visium HD FFPE and Visium HD FF data were processed with spaceranger (v.3.0.0). Stereo-seq v1.3 data were processed with SAW (v.8.0). GRCh38 was used as the reference genome. For Xenium 5K, we retained the calls with Phred-scaled quality higher than 20. The calls from CosMx 6K underwent filtration based on the methodologies described in the previous study^22^. The DAPI images of all FOVs from CosMx 6K were stitched using napari-cosmx (https://github.com/Nanostring-Biostats/CosMx-Analysis-Scratch-Space). We performed tissue cut to remove the calls outside the tissue using the Python package OpenCV (v.4.10.0) for Xenium 5K and CosMx 6K. For a fair comparison, we binned Stereo-seq v1.3, Xenium 5K, and CosMx 6K data at the resolution of 8 μm and used the 8-μm output of Visium HD FF and Visium HD FFPE for basic metric evaluations.

### Correlation for gene expression levels between ST and scRNA-seq data

The total transcript count of each gene was log10 transformed, and the correlation of gene expression levels between ST data and scRNA-seq data was calculated. For CosMx 6K, calls were ranked based on their probability of being random signals. We then stepwise extracted the top percentages of high-quality calls, specifically at 50%, 62.5%, 75%, and 87.5%, and assessed their correlation with scRNA-seq data.

### Downsample sequencing reads for Stereo-seq v1.3 and Visium HD FFPE data

The –-unmapped-fastq option was used in SAW to retain unmapped reads for Stereo-seq v1.3. We selected ten regions characterized by a high density of cancer cells based on H&E staining for each dataset. BAM files were filtered to isolate the reads with valid UMI information. The valid reads with spatial coordinates mapped to the ten regions were downsampled at fixed proportions (20%, 40%, 60%, and 80%) with the Python package pysam (v.0.22.1).

### Evaluation of diffusion for Stereo-seq v1.3 and Visium HD FFPE data

The transcript count of each bin outside the tissue region was normalized by the mean transcript count inside the tissue region. We defined the diffusion distance as the distance between each bin outside the tissue region with its nearest neighbor bin located within the tissue region, which was obtained with the NearestNeighbors function in Python package scikit-learn (v.1.5.2). Given the larger chip size of the Stereo-seq v1.3 and its potential for long-range transcript diffusion, we only retained bins with distances to tissue shorter than the maximum diffusion distances observed for Visium HD FFPE.

### Registration of images and alignment of spatial data

We manually annotated key landmarks on each image and employed the SimpleITK library (v.2.4.0) to achieve accurate registration of paired images. Specifically, the Similarity2DTransform, SetMetricAsMattesMutualInformation, and sitkLinear functions were utilized to perform the automatic adjustment after the initial transform obtained with paired landmarks.

For the alignment of Visium HD FFPE, CosMx 6K, and Xenium 5K data, we set the grayscale H&E image of Visium HD FFPE as the fixed reference and registered the DAPI images of CosMx 6K and Xenium 5K to it. The derived transformations were subsequently applied to map the CosMx 6K and Xenium 5K data onto the coordinate system of Visium HD FFPE data. For the alignment of ST data with adjacent CODEX data, the grayscale H&E images of Stereo-seq v1.3, Visium HD FFPE, and Visium HD FF were used as the fixed references. For CosMx 6K and Xenium 5K, the DAPI images were used as the fixed references. The fixed references were rescaled to match the resolution of CODEX, and the DAPI channel of CODEX was registered to them. The derived transformations were subsequently applied to the remaining CODEX channels. To enable direct comparisons across FFPE samples, we used the Python package OpenCV to extract tissue masks of each ST data based on the paired staining images and intersected them to define the shared regions. A similar approach was used to extract overlapping regions between ST data and adjacent CODEX data.

### Annotation of scRNA-seq data

Genes detected in fewer than 10 cells were excluded from the analysis. Cells that did not fulfil the following criteria were removed:

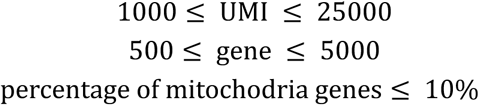

Putative doublets were identified and removed using DoubletFinder^52^ (v.2.0.3). A two-round clustering strategy was applied for cell type annotation using Seurat^53^. In the first round of clustering, the data were normalized and log-transformed to the same scale. A set of 2,000 highly variable genes was identified, followed by scaling of the expression matrix. The top 30 principal components (PCs) were identified to build nearest-neighbor graph. Clustering was performed using the shared nearest neighbor (SNN) modularity optimization algorithm. We annotated each cluster based on its expression of the following known markers: B cell, *CD79A*, *CD19*, and *MS4A1*; cDC1, *XCR1* and *CLEC9A*; cDC2, *CD1C* and *CLEC10A*; mregDC, *LAMP3* and *CCR7*; pDC, *LILRA4*; macrophage, *CD68*, *C1QC*, and *SPP1*; mast cell, *KIT* and *TPSAB1*; monocyte, *FCN1*; neutrophil, *CSF3R* and *AQP9*; endothelial, *VWF*, *CD34*, *CDH5*, and *PECAM1*; fibroblast, *ACTA2*, *COL1A2*, and *FAP*; SMC, *ACTA2* and *RGS5*; NK cell, *FCGR3A*, *GZMA*, and *NCAM1*; plasma cell, *SDC1* and *MZB1*; CD4+ T cell, *CD4*, *CD3G*, *CD3D*, and *CD3E*; CD8+ T cell, *CD8A*, *CD8B*, *CD3G*, *CD3D*, and *CD3E;* Tprolif, *MKI67*, *CD3G*, *CD3D*, and *CD3E*; epithelial*, EPCAM*; hepatocyte, *ALB*; kupffer cell, *CD5L*. The subtypes of T cells were annotated after a second round of clustering using a similar approach.

### Evaluation of segmentation results

Solidity is calculated as the ratio of the contour area to its convex hull area, with values approaching 1 indicating convexity and lower values suggesting concavity. Circularity measures how closely a shape resembles an ideal circle, with 1 corresponding to a perfect circle and 0 indicating a more irregular shape. The aspect ratio is the ratio of the width to the height of the bounding box that encloses the contour, which describes the elongation of the shape, with values greater than 1 indicating horizontal elongation and values less than 1 indicating vertical elongation. All values were calculated using the Python library OpenCV.

### Evaluation of the clustering of ST data based on the transcriptomics profile

Genes detected in fewer than 100 bins or cells were filtered out. Bin-level data demonstrated lower quality than cell-level data due to the retention of non-cellular regions that would be excluded after cell segmentation. To address this, we applied a more stringent cutoff to filter out the low-quality bins. For Visium HD FFPE, bins with total counts below the 20th percentile of all bins were excluded. For cell-level data from Stereo-seq v1.3, CosMx 6K, and Xenium 5K, we filtered out low-quality cells with total counts below the 10th percentile of all cells. We further utilized the Python package scanpy (v.1.10.3) to perform clustering. Data were normalized and log-transformed to the same scale. The top 10% of genes with the highest variance were defined as high-variable genes. The top 30 principal components were computed to build neighborhood graphs. Data were embedded using Uniform Manifold Approximation and Projection (UMAP) for further dimensionality reduction and visualization. Clustering was performed using the Leiden algorithm with default resolution settings. The silhouette_score function from the Python package scikit-learn was used to assess the clustering quality. This score evaluates cluster separation by comparing intra-cluster and inter-cluster distances, with a value approaching 1 indicating better-defined clusters.

### Annotation of ST and CODEX data

The latest versions of SELINA^37^, Celltypist^38^, Spoint^39^, Tangram^40^, and TACCO^41^ were used to transfer the annotations from scRNA-seq data to the filtered ST data. We utilized two metrics to evaluate the annotation consistency across different tools: (1) the proportion of cells that were consistently annotated as the same cell type by different tools; (2) the entropy of cell numbers detected by different tools for each cell type, which was calculated with the following formula:

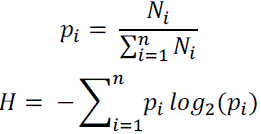

where the cell number detected by the i-th tool was denoted with *N*_*i*_ and its ratio over all tools was denoted with *p*_*i*_. We further integrated the annotations from these five tools using a majority-vote approach. Each cell was assigned with the cell type supported by most tools. For cells with annotations differing across tools, the method demonstrating the highest alignment with other tools was applied to resolve discrepancies and assign the final cell type annotation.

CODEX data exhibited prominent non-specific binding signals in tumour regions and background signals across whole sections, which could bias cell annotation if based solely on the average signal intensity. To address this challenge, we manually labeled hundreds of positive cells for each marker and trained a K-nearest neighbor (KNN) classifier in QuPath. For membrane markers, truly positive cells were defined as those with fluorescent signals surrounding the cell nuclei. For transcription factors, positive cells were those with fluorescent signals confined exclusively to the nucleus. Cells lacking any marker signal were categorized as negative cases. We trained the classifier with various signal statistics, including mean, median, minimum, maximum, and standard deviation for signals in the nucleus, cytoplasm, membrane, and the entire cell. This classifier was then applied to annotate the remaining cells across the whole section.

### Correlation between ST and CODEX data

For the correlation analyses between ST transcripts and CODEX pixel density, CODEX channels with strong nonspecific binding or low abundance were excluded. To further mitigate the influence of CODEX background signals, cell segmentation was performed on the DAPI channel using Stardist^54^ (v.0.5.0) and QuPath^55^ (v.0.5.1). Only protein staining signals confined within the segmented cells were retained for the subsequent analysis. We binned the shared regions of ST and CODEX data at multiple spatial resolutions (100, 200, 300, 400, and 500 μm). For each bin, pixel density and transcript counts were aggregated. Besides, the top 50 differentially expressed genes for each cell type obtained with paired scRNA-seq data were aggregated as the cell-type signature. Correlations between transcript counts or cell-type signatures and CODEX pixel density were calculated over the spatial grids. Furthermore, the correlation of cell distribution between ST data and CODEX data was determined using a similar approach for each cell type.

### Spatial clustering of ST and CODEX data

CellCharter^42^ was used to perform spatial clustering on both ST and CODEX data. CODEX can accurately measure the spatial distribution of marker proteins across major cell types, offering insights into the underlying tissue architecture. Given this, we leveraged the optimal cluster number derived from CODEX data to guide the clustering of ST data. To investigate the alignment of clustering results between CODEX and ST data, we computed cell proportions within spatial grids (100 × 100 μm) for each cluster and assessed their correlations. The correlation coefficient is defined as:

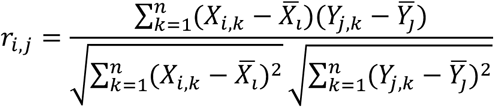

where *r_i,j_* is the correlation coefficients matrix, *X_i,k_* and *Y_j,k_* are the proportions of cluster *i* in ST data and cluster *j* in CODEX data for the *k*-th bin, and 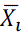 and 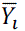 are the respective means. We applied the Hungarian algorithm to identify the optimal cluster pairs between CODEX and ST data, aiming to maximize the correlation between the two modalities. The consistency was finally quantified as the average correlation across all paired clusters.

## Data availability

The raw and processed datasets are publicly accessible on the SPATCH website at http://spatch.pku-genomics.org/. Beyond downloading, this web server offers tools for data visualization and exploration, enabling users to interactively analyze the datasets.

## Code availability

The code utilized for data processing and analysis in this study is publicly available on GitHub (https://github.com/zenglab-pku/SPATCH).

## Acknowledgments

This work was supported by the National Natural Science Foundation of China (92374116, 82341026, 32470664, and 12226005) and Peking-Tsinghua Center for Life Sciences. We thank the Optical Imaging Core Facility, National Center for Protein Sciences at Peking University in Beijing, China, for assistance with the CODEX experiment.

## Author contributions

Z.X.Z., Z.Z. and Y.M. conceived the study and designed experiments. P.R. performed the bioinformatics analysis with help from Y.W., C.L., and S.W. R.Z., Y.L. and X.L. collected and processed the tumour samples. P.Z. developed the SPATCH website. Zongxu Zhang, Y.Z., and Y.H. helped with the study design and analysis of ST data. P.R., R.Z., Z.Z. and Z.X.Z. wrote and revised the manuscript with the help of other authors.

## Competing interests

The authors declare no competing interests.

**Supplementary Fig. 1.**
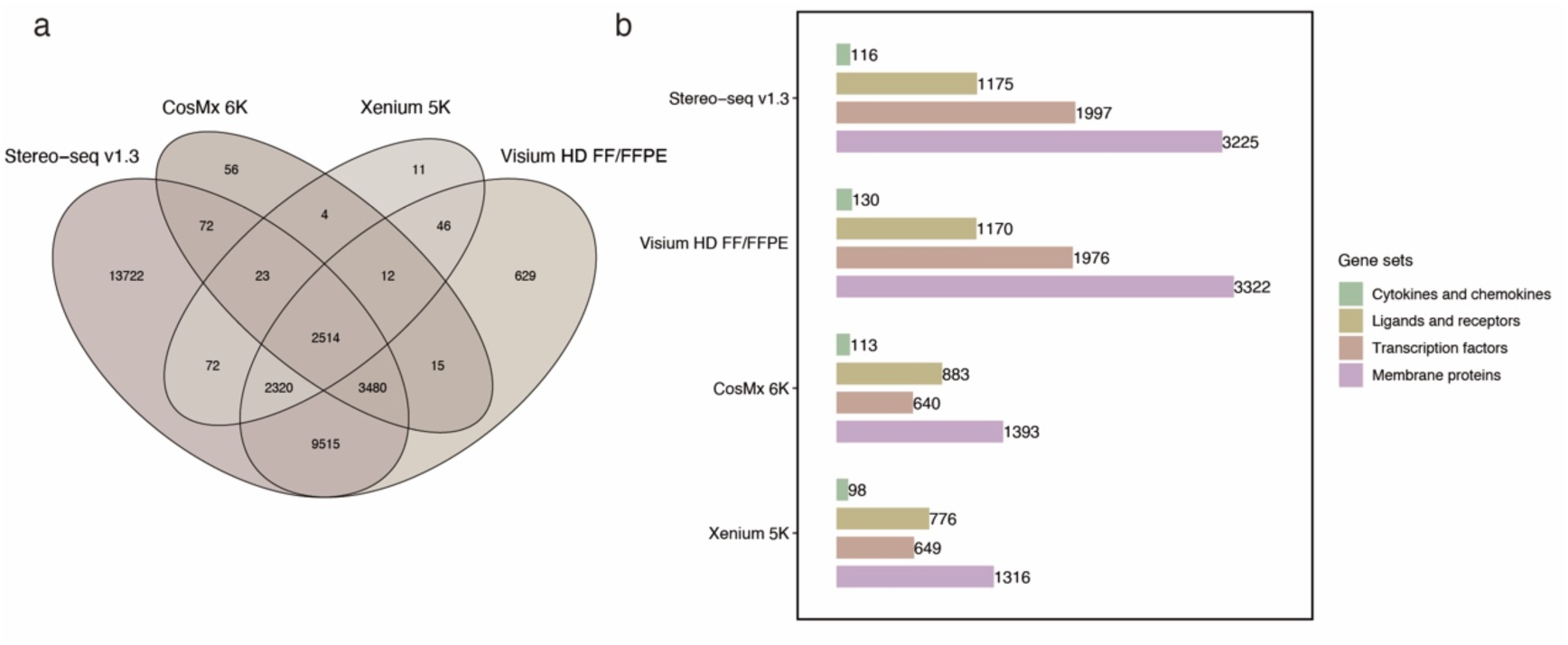
Comparison of genes detected by different ST platforms. **a.** Overlap of genes detected by different ST platforms. **b.** Overlap between genes detected by different ST platforms and functional gene sets, including cytokines, chemokines, ligands, receptors, transcription factors, and membrane proteins.

**Supplementary Fig. 2.**
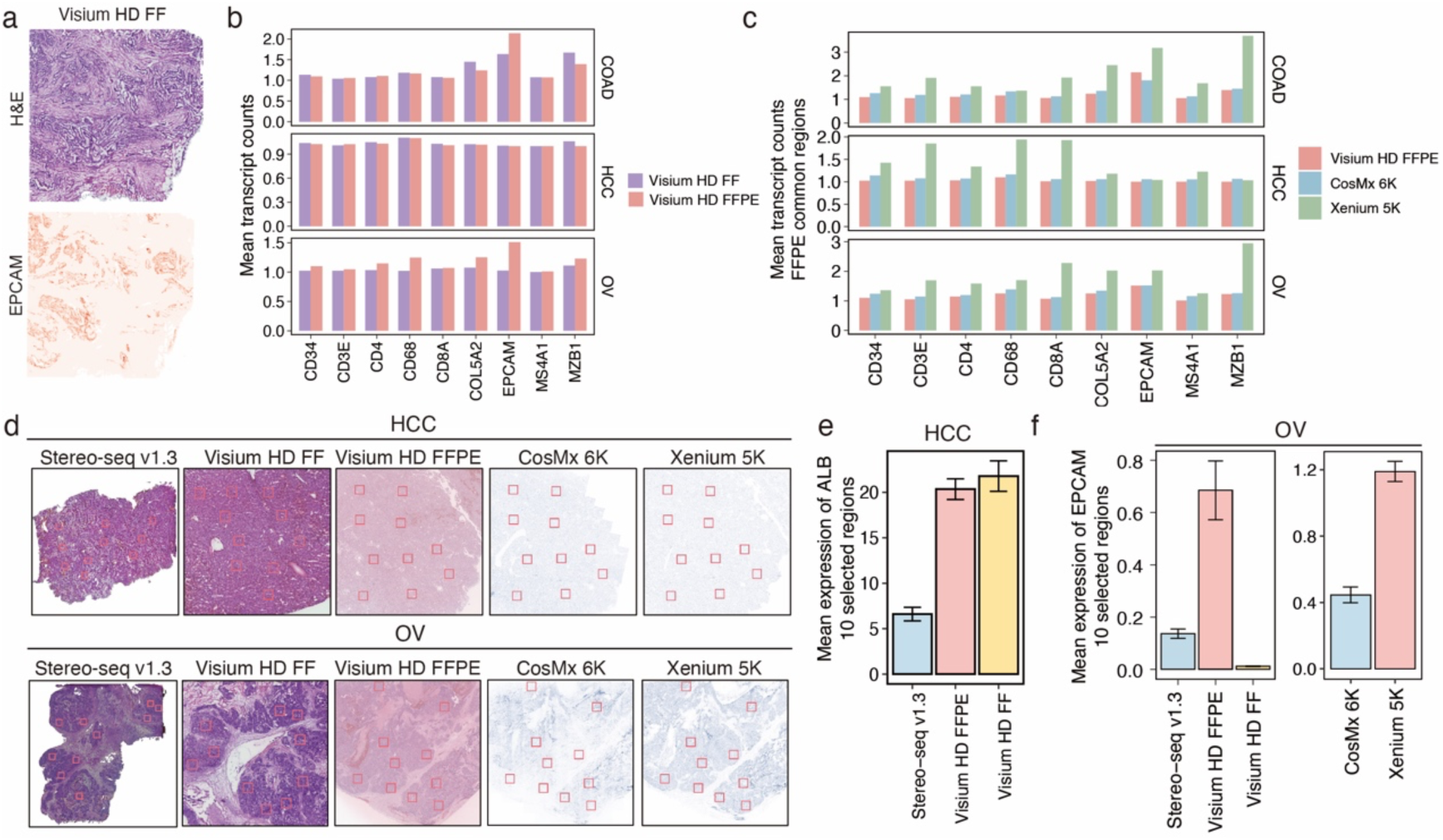
Comparison of sensitivity for lineage marker genes. **a.** H&E staining (top) and spatial distribution of EPCAM transcripts (bottom) in the COAD section profiled with Visium HD FF. Color intensity represents transcript count in each 8 × 8 μm bin. **b.** Mean expression levels of lineage marker genes across whole sections for Visium HD FFPE and Visium HD FF in three cancer types at 8-μm resolution. **c.** Mean expression levels of lineage marker genes within the shared regions of Visium HD FFPE, CosMx 6K, and Xenium 5K data at 8-μm resolution**. d.** Spatial distribution of the ten selected regions (400 × 400 μm for each one) with similar morphology in HCC and OV, highlighted by red squares. **e-f.** Average transcript counts of ALB (hepatocyte marker, **e**) and EPCAM (epithelial marker, **f**) for all 8 × 8 μm bins within the ten selected regions in HCC (**e**) and OV (**f**). Error bars indicate the SEM across the ten selected regions.

**Supplementary Fig. 3.**
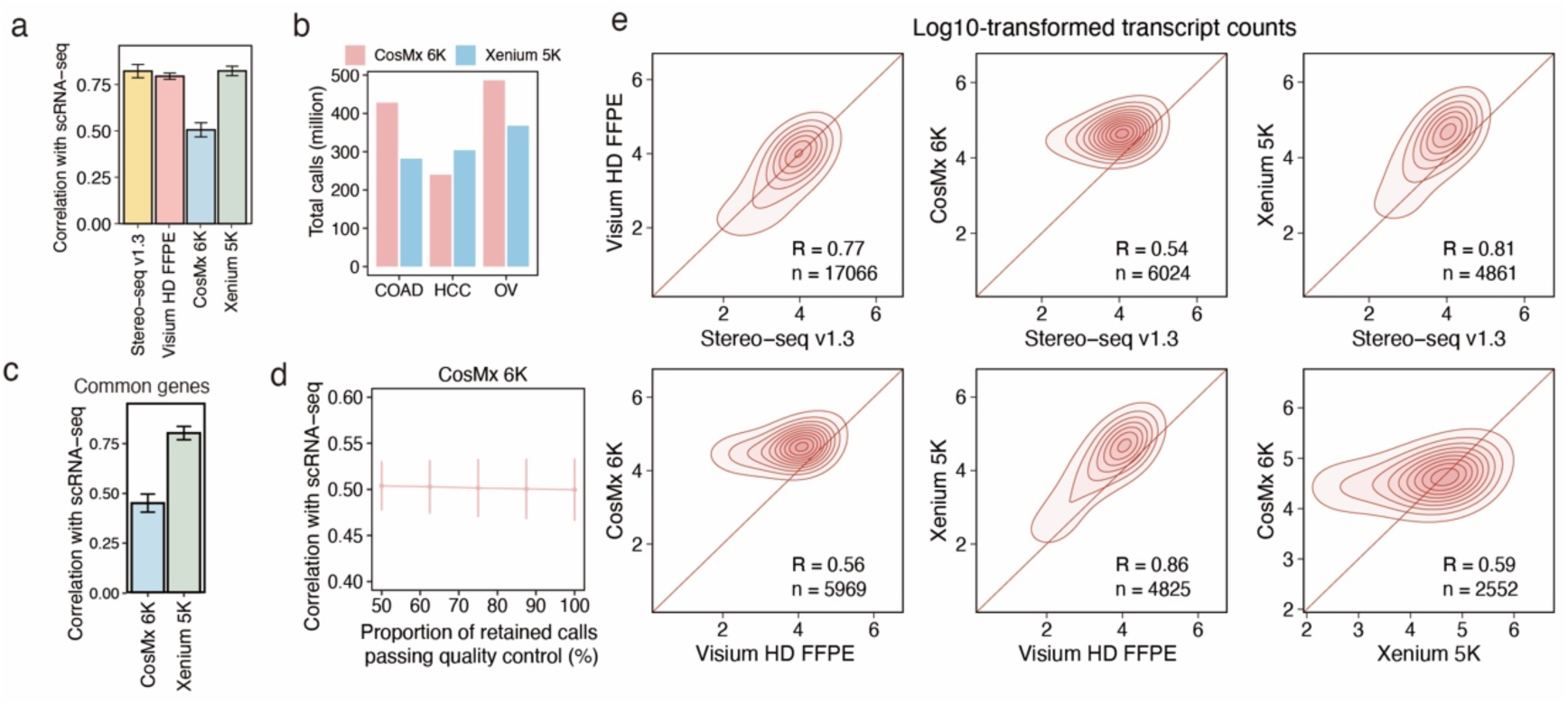
Correlation of gene expression levels across diverse platforms. **a.** Pearson correlation of gene expression levels between ST and scRNA-seq data. SEM across three cancer types is shown. **b.** Total number of calls detected by CosMx 6K and Xenium 5K. **c.** Pearson correlation of gene expression levels between CosMx 6K, Xenium 5K, and scRNA-seq data. Shared genes (n = 2552) between CosMx 6K and Xenium 5K data were used in the analysis. SEM across three cancer types is shown. **d.** Pearson correlation of transcript counts between CosMx 6K and scRNA-seq data, calculated at stepwise decreasing thresholds to filter the high-quality calls. SEM across three cancer types is shown. **e**. Paired comparison of gene expression levels across different ST platforms. For each gene, the total transcript counts across three cancer types were averaged and log10 transformed. The diagonal red line indicates a slope of 1, and color intensity corresponds to relative gene counts. R denotes the correlation coefficient, and n indicates the number of genes included in the analysis.

**Supplementary Fig. 4.**
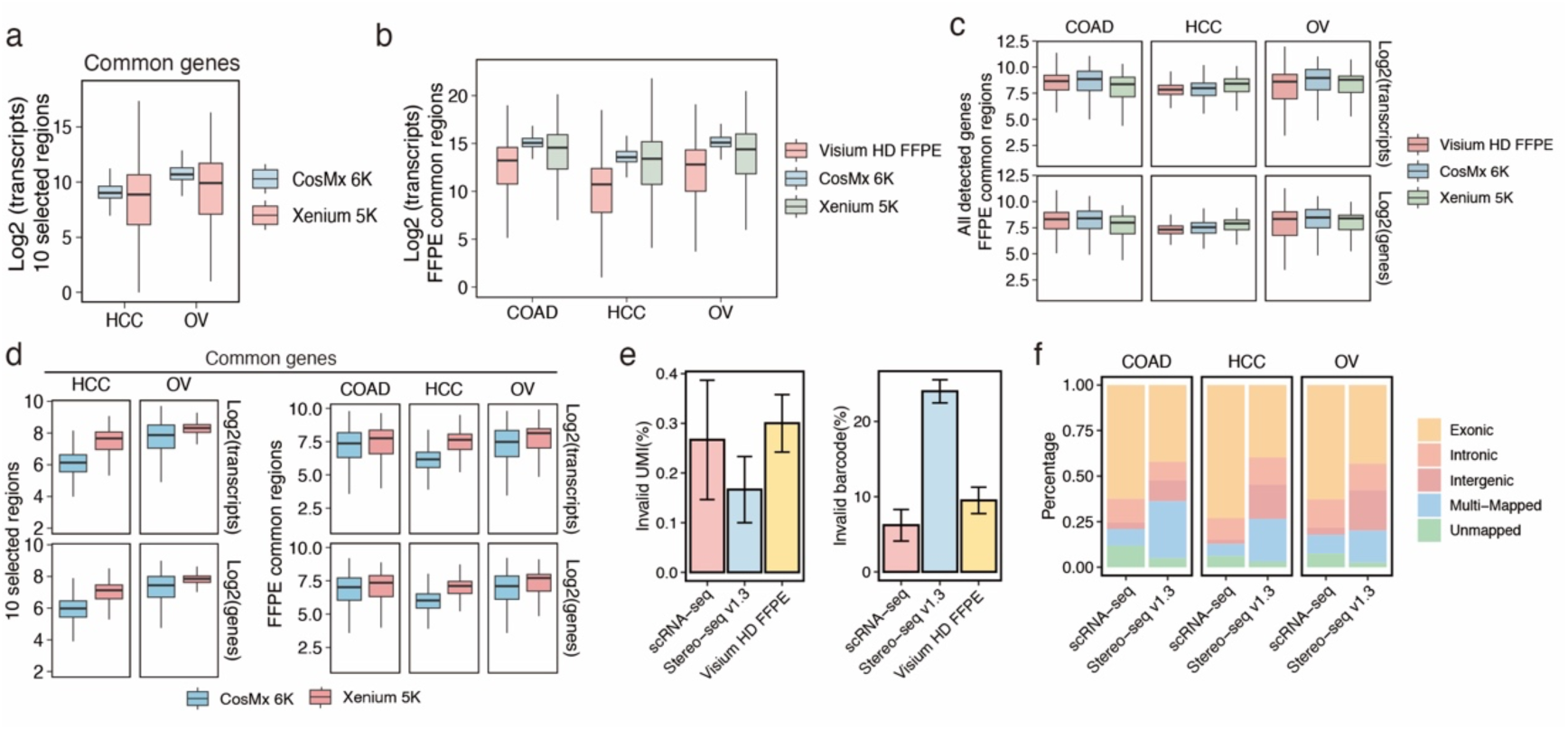
Comparison of sensitivity for all genes detected by each ST platform. **a.** Log_2_-transformed transcript count for shared genes (n = 2552) between CosMx 6K and Xenium 5K data in the ten selected regions. **b.** Log_2_-transformed transcript count for each gene within the shared regions of Visium HD FFPE, CosMx 6K, and Xenium 5K data. **c.** Log_2_-transformed transcript count (top) and gene count (bottom) for each 8-μm bin within the common regions of Visium HD FFPE, CosMx 6K, and Xenium 5K data. All detected genes were included in the analysis. **d.** Log_2_-transformed transcript count (top) and gene count (bottom) for each 8-μm bin within the ten selected regions (left) or common regions (right) between CosMx 6K and Xenium 5K data. Common genes (n = 2552) between CosMx 6K and Xenium 5K data were included in the analysis. **e.** Percentage of reads failing quality control for UMIs (left) and reads with barcodes not matching the whitelist or mask file (right). SEM across three cancer types is shown. **f.** Proportion of reads mapped to different genomic regions: exonic, intronic, intergenic, multi-mapped, and unmapped in the human genome.

**Supplementary Fig. 5.**
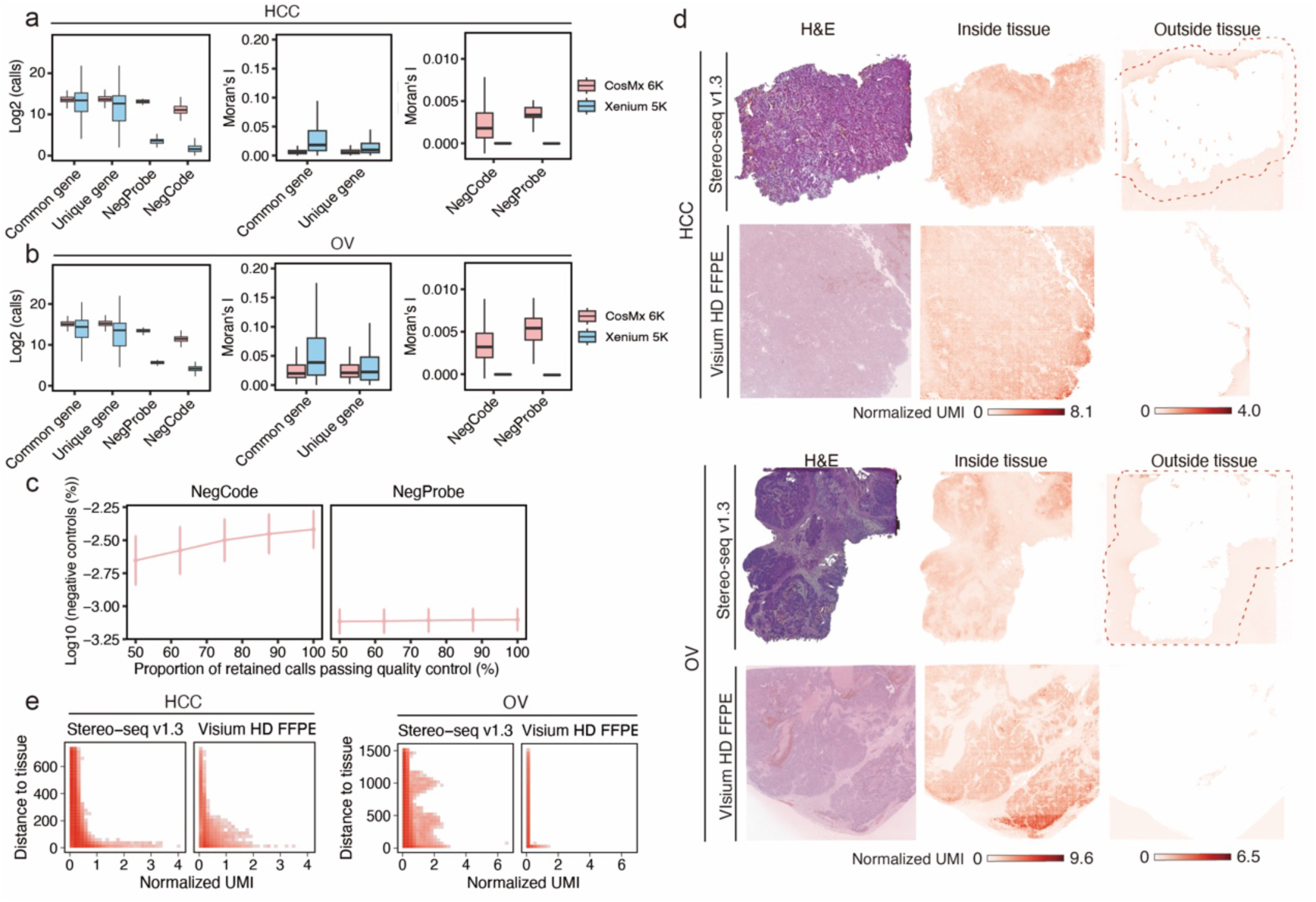
Evaluation of negative control signals and transcript diffusion. **a-b.** Number of calls and Moran’s I (spatial autocorrelation) for common genes, platform-specific genes (unique gene), negative control sequences (NegProbe), and negative control codes (NegCode) detected by CosMx 6K and Xenium 5K in HCC (**a**) and OV (**b**). **c.** Percentage of negative control signals for CosMx 6K data, calculated at stepwise decreasing thresholds to filter the high-quality calls. SEM across three cancer types is shown. **d.** H&E staining (left), spatial distribution of transcripts detected inside (middle) and outside (right) the tissue region for HCC and OV. Regions of Stereo-seq v1.3 data used for downstream analysis are highlighted with red dashed lines. Color intensity indicates the mean-normalized transcript count in each 8 × 8 μm bin. **e.** Diffusion evaluation in HCC (left) and OV (right). The x-axis represents the mean-normalized transcript count for each bin outside the tissue. The y-axis shows the minimum distance from bins outside the tissue region to the tissue boundary. Color intensity indicates the number of bins.

**Supplementary Fig. 6.**
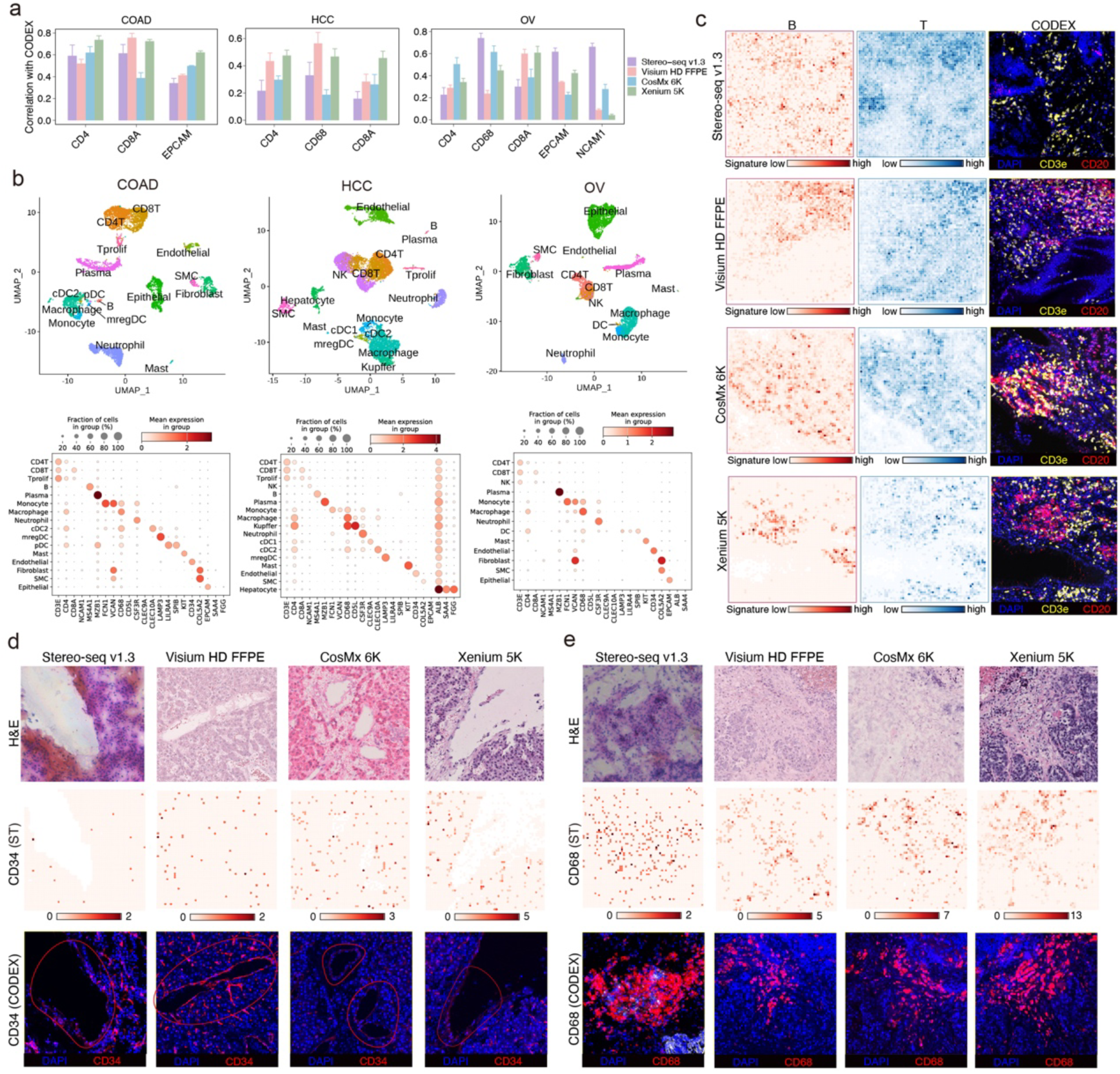
Spatial correlation of transcriptomic and proteomic data across key cell types and tissue structures. **a.** Spatial correlation between transcript counts and the corresponding protein abundance over the spatial grids. Pearson correlation coefficients are reported. Error bars represent SEM for correlations obtained under different grid sizes. **b.** UMAP representation of scRNA-seq data from three cancer types (top) and expression levels and percentage of lineage marker genes in different cell types (bottom). **c.** Spatial distribution of B cell (left) and T cell (middle) signatures, along with staining of corresponding marker proteins (right), within ROIs (500 × 500 μm) showing high lymphocyte infiltration in COAD. Color intensity represents the signature score in each 8 × 8 μm bin. **d.** H&E staining (top), spatial distribution of CD34 transcripts (middle), and CD34 protein staining (bottom) within the ROIs (500 × 500 μm) containing the vascular structures that are highlighted with red solid lines in HCC. Color intensity represents the transcript count in each 8 × 8 μm bin. **e.** H&E staining (top), spatial distribution of CD68 transcripts (middle), and CD68 protein staining (bottom) within the ROIs (500 × 500 μm) containing the macrophage aggregates in OV. Color intensity represents the transcript count in each 8 × 8 μm bin.

**Supplementary Fig. 7.**
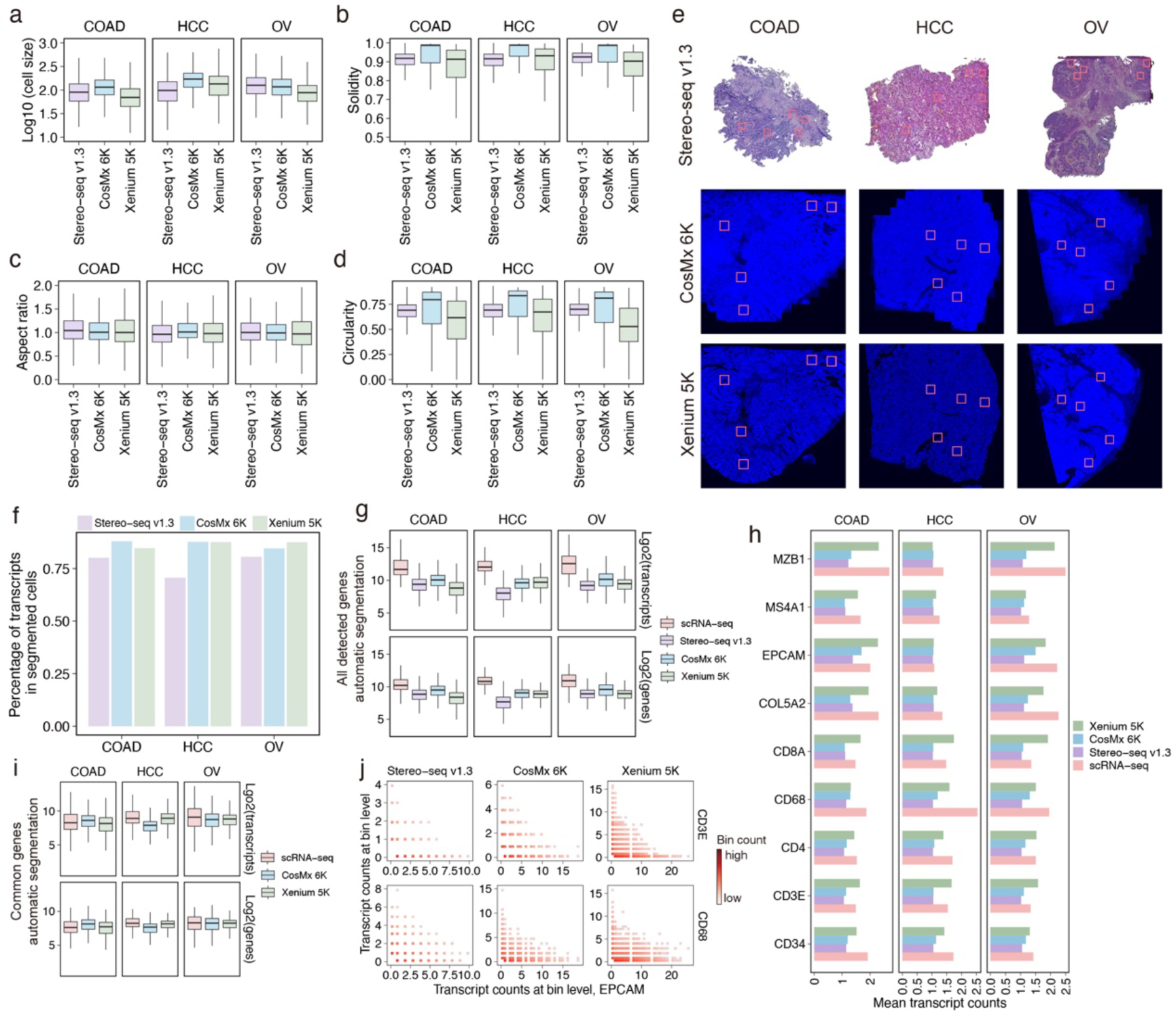
Comparison of cell segmentation. **a-d.** Quantitative features of automatically segmented cells, including cell size **(a)**, solidity **(b)**, aspect ratio **(c)**, and circularity **(d)** for each platform. **e.** Spatial distribution of regions (500 × 500 μm) selected for comparing manually annotated cell boundaries with automatic segmentations across different ST platforms. **f.** Percentage of transcripts within the automatically segmented cells. **g.** Log_2_-transformed transcript and gene counts of each automatically segmented cell for all detected genes. **h.** Mean expression levels of lineage marker genes for all cells from scRNA-seq data and automatical segmentation results from ST data. **i.** Log_2_-transformed transcript and gene counts of each automatically segmented cell for common genes detected by the two iST platforms and scRNA-seq. **j.** Density plot of CD3E/CD68 and EPCAM expressions at bin level in COAD. Only bins containing at least one transcript of either CD3E/CD68 or EPCAM were included. Color intensity indicates the number of 8× 8 μm bins.

**Supplementary Fig. 8.**
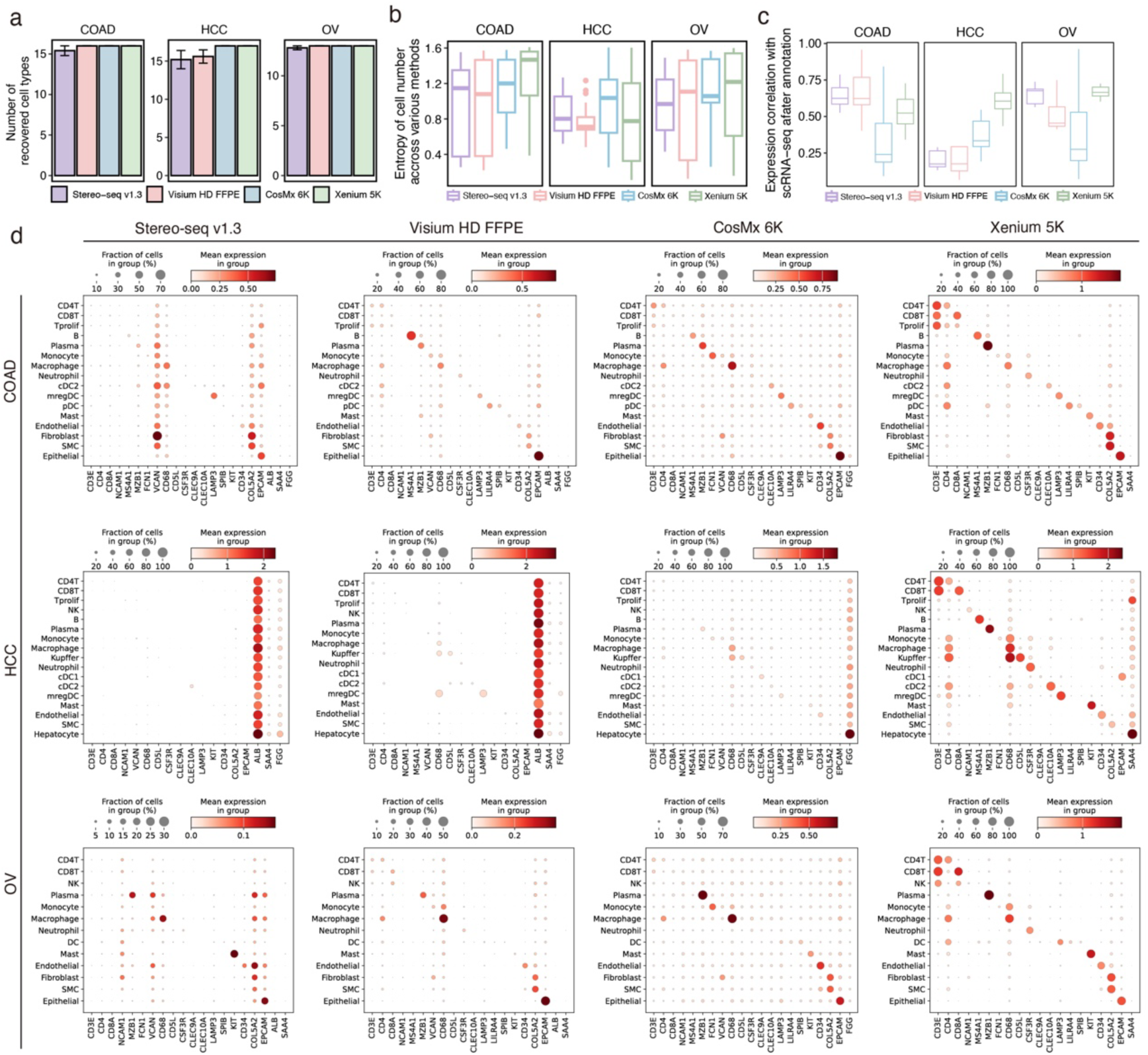
Consistency between ST and scRNA-seq data in terms of cell type identification. **a.** Number of annotated cell types recovered by different ST platforms. **b.** Entropy of cell numbers obtained using various annotation tools for each cell type. **c**. Pearson correlation of gene expression between ST data and scRNA-seq data for each annotated cell type. **d**. Expression levels and percentage of lineage marker genes in different cell types.

**Supplementary Fig. 9.**
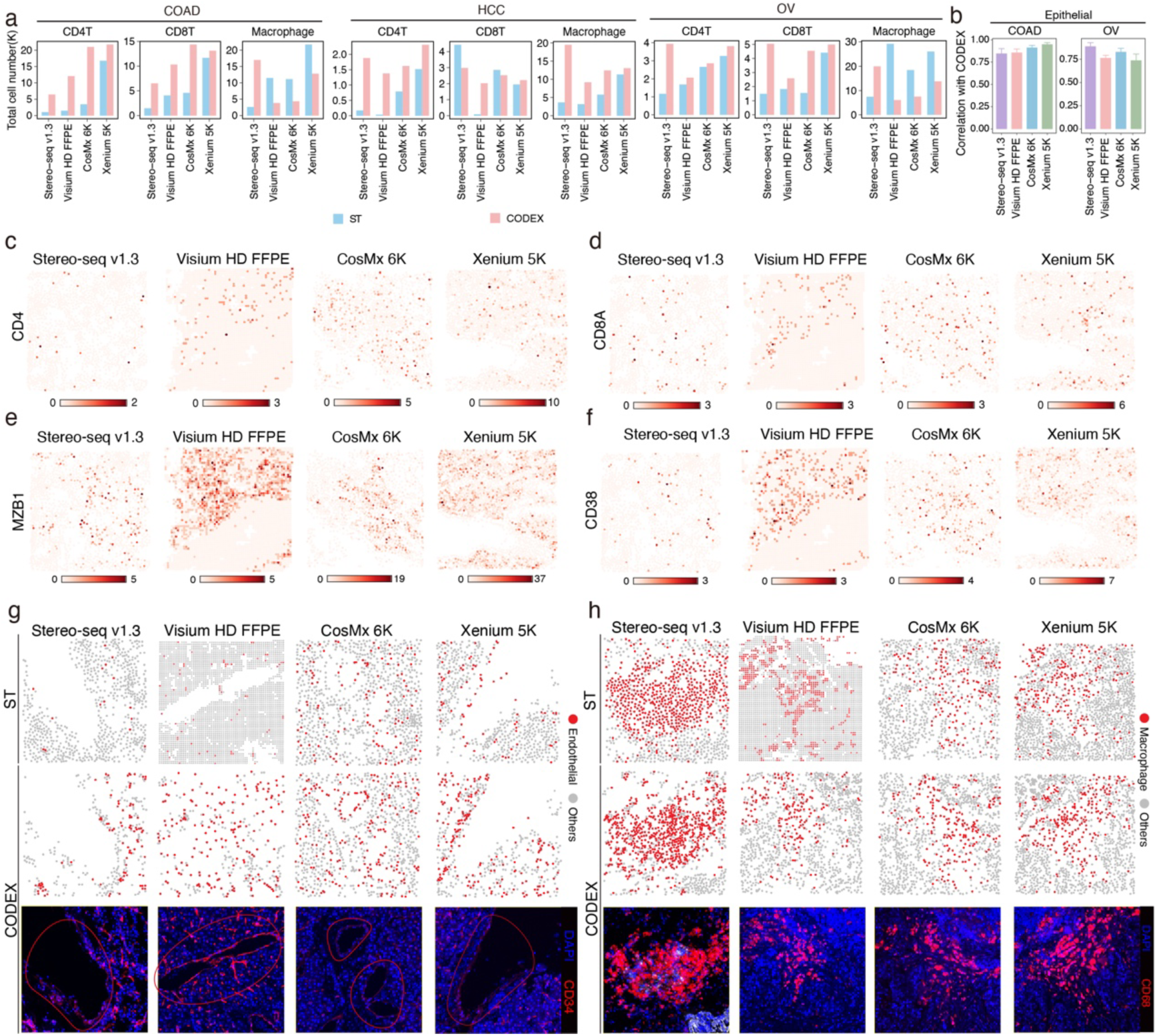
Characterization of spatial distribution for major cell types using ST data and CODEX data. **a.** Total number of immune cells detected by different ST platforms and corresponding adjacent CODEX across entire sections. **b.** Correlation of cell counts between ST data and adjacent CODEX for epithelial cells over spatial grids in COAD and OV. Pearson correlation coefficients are shown. Error bars represent SEM for correlations obtained under different grid sizes. **c-f.** Spatial distribution of CD4 (**c**), CD8A (**d**), MZB1 (**e**), and CD38 (**f**) transcripts within ROIs (500 × 500 μm) showing high lymphocyte infiltration. **g-h.** Spatial distribution of endothelial cells within ROIs (500 × 500 μm) containing vascular structure in HCC (**g**) and macrophages within ROIs (500 × 500 μm) containing macrophage aggregates in OV (**h**). The top row shows different ST data, the middle row shows annotations from adjacent CODEX, and the bottom row shows the staining for CD34 (**g**) and CD68 (**h**).

**Supplementary Fig. 10.**
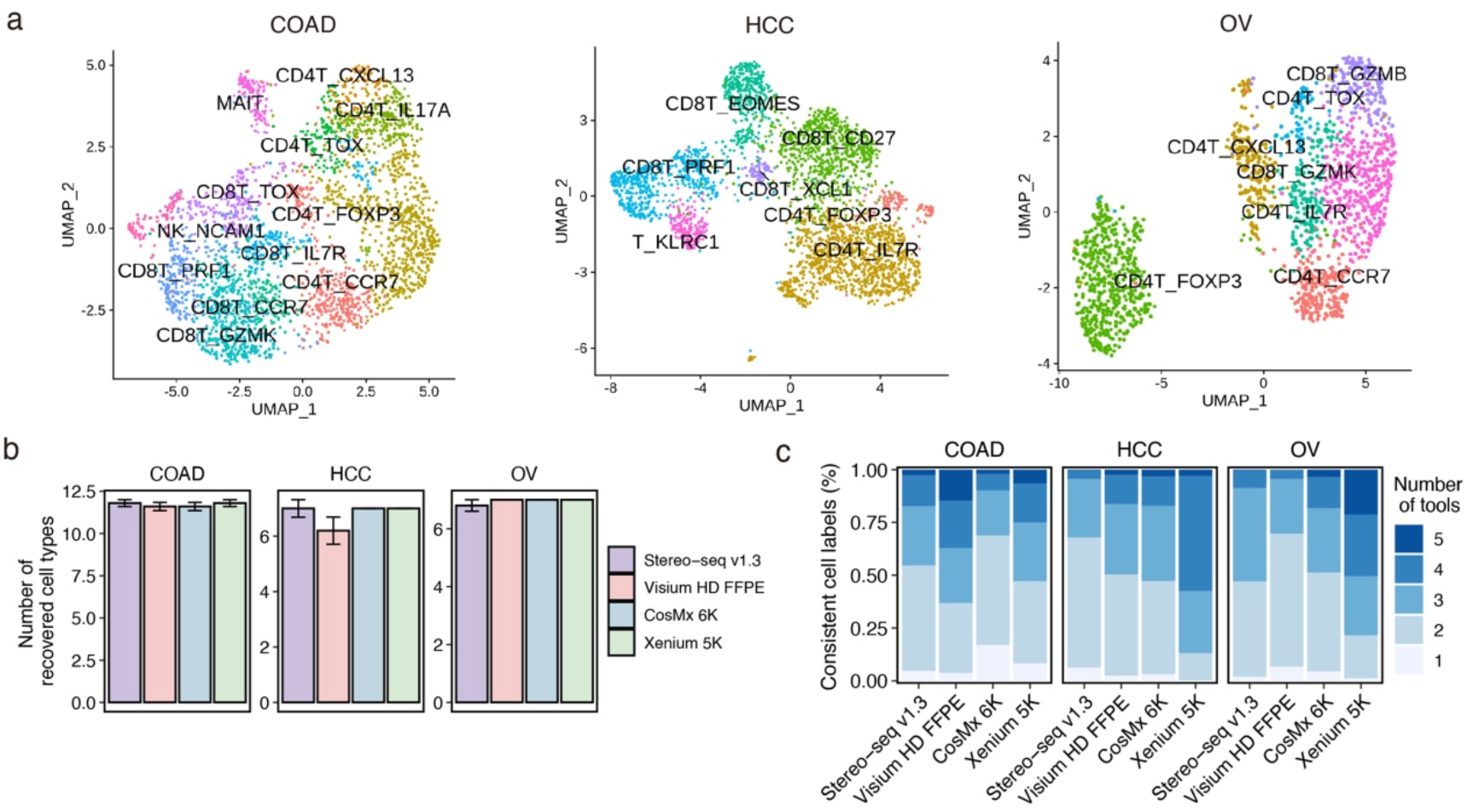
Characterization of T cell subtypes. **a.** UMAP representation of scRNA-seq data from three cancer types, colored by T cell subtypes. **b.** Number of T cell subtypes recovered by different ST platforms. **c.** Percentage of T cells consistently annotated as the same subtype by multiple automatic annotation tools.

**Supplementary Fig. 11.**
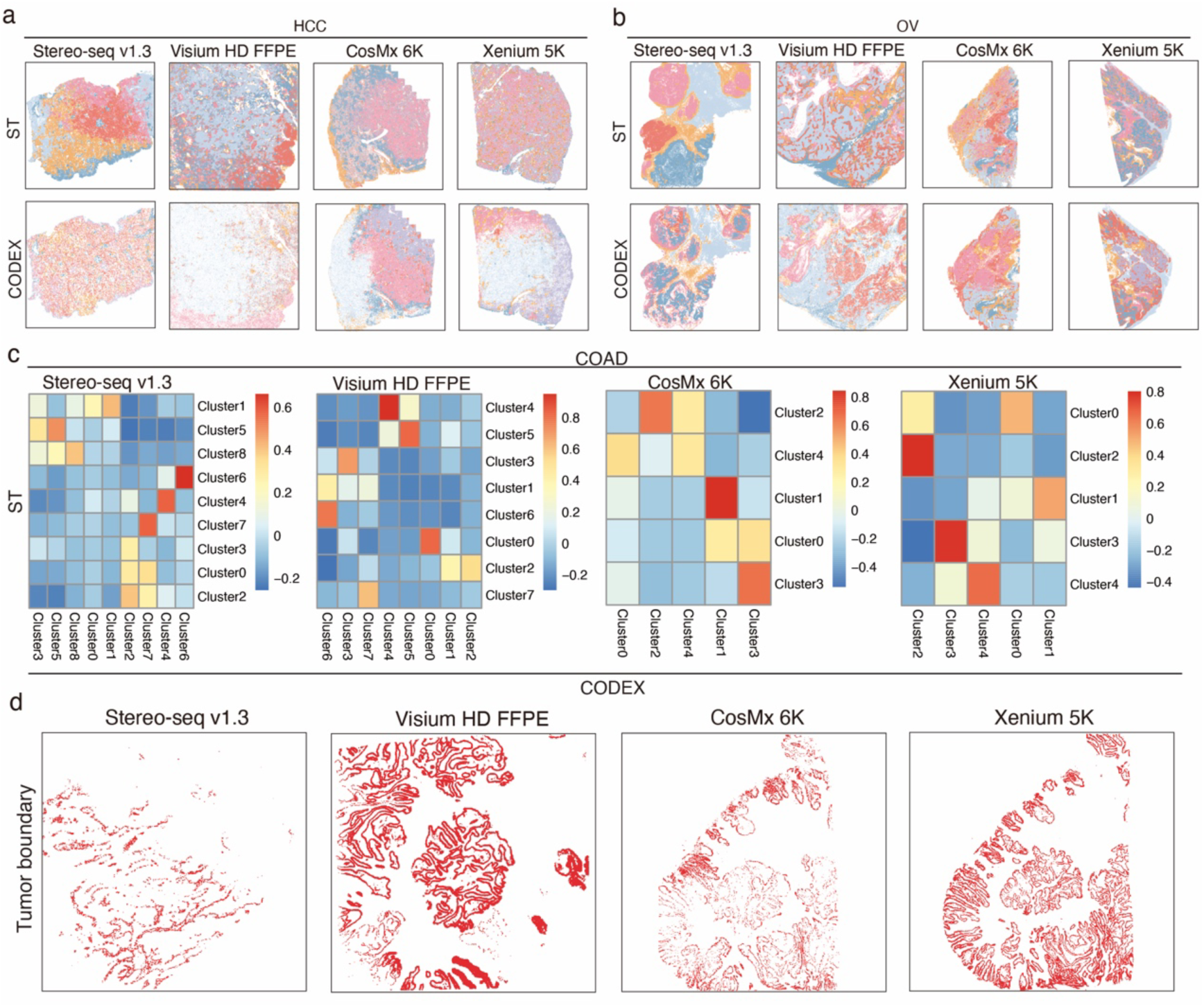
Spatial clustering and malignant cell localization. **a-b.** Spatial clustering of ST (top) and CODEX data (bottom) in HCC (**a**) and OV (**b**), with distinct colors representing different spatial clusters. **c.** Correlation between the proportions of spatial clusters identified from CODEX and ST data across all spatial grids (100 × 100 μm). Rows correspond to the spatial clusters from ST data; columns correspond to the spatial clusters from CODEX; color intensity denotes the degree of correlation. **d.** Spatial distribution of malignant cells localized at the tumour boundary.

**Supplementary Table 1 Timeline for sample collection and processing**

**Supplementary Table 2 Technical information for all ST platforms used in this study**

**Supplementary Table 3 Gene sets collected from previous studies**

**Supplementary Table 4 Intersection between reference gene sets and gene panels of each ST platform**

**Supplementary Table 5 Marker genes used in the mutually exclusive expression analysis**

**Supplementary Table 6 Antibodies and cycle information for the CODEX experiment**

